# Cytopath: Simulation-based inference of differentiation trajectories from RNA velocity fields

**DOI:** 10.1101/2020.12.21.423801

**Authors:** R. Gupta, D. Cerletti, G. Gut, A. Oxenius, M. Claassen

## Abstract

Trajectory inference from single-cell RNA sequencing data bears the potential to systematically reconstruct complex differentiation processes, however inferring trajectories that accurately model the biological characteristics of varied processes continues to be a challenge, notwithstanding the many available solutions. In general, trajectory and pseudotime inference methods have so far suffered from the ambiguity of static single-cell transcriptome snapshots lacking a concept of directionality and rate of transcriptional activity.

We report Cytopath, a method for trajectory inference that takes advantage of transcriptional activity information from RNA velocity of single-cells to perform trajectory inference. Cytopath performs this task by defining a Markov chain model, simulating an ensemble of possible differentiation trajectories and constructs a consensus trajectory. We show that Cytopath can recapitulate the topological and molecular characteristics of the differentiation process under study. In our analysis we include differentiation trajectories with varying bifurcated, circular, convergent and mixed topology studied in single-snapshot as well as time-series single-cell RNA sequencing experiments. We demonstrate superior and enabling capability to reconstruct differentiation trajectories in comparison to state-of-the art trajectory inference approaches.

## 1 Introduction

Biological processes such as cell type differentiation [Manno et al., 2018, Bastidas-Ponce et al., 2019, Burns et al., 2015], immune response [Cerletti et al., 2020] or cell division [Mahdessian et al., 2021] can be conceptualized as temporal sequences of coordinated, phenotypic state changes in the context of, possibly heterogeneous, cell populations. Such phenotypic states can be characterized by e.g. epigenetic, transcriptional and proteomic cell profiles. Furthermore, these differentiation processes are often asynchronously triggered. The differentiation processes give rise to state sequences with varying topologies including bi-furcating, multi-furcating, cyclical and convergent trajectories.

This situation requires single-cell approaches to measure and ultimately investigate these processes. The repertoire of suitable technologies to monitor different types of molecular profiles has increased dramatically over the last years. In particular single-cell RNA sequencing (scRNAseq) has gained wide-spread use due to the broad applicability of sequencing technology. While these measurements are information rich, their analysis and interpretation is challenged by high dimensionality, low sequencing depth, measurement noise and its destructive nature only yielding snapshots of the whole process.

Different computational approaches have been proposed to model differentiation processes from scRNAseq data, specifically covering the tasks of pseudotime estimation, trajectory inference or cell fate prediction. These tasks are related but typically require different approaches (Table S1). The goal of cell fate prediction is to determine the terminal differentiation state (fate) of any cell, possibly already early in the differentiation process. Such methods generate a score or probability per cell with respect to terminal differentiation states [Lange et al., 2022, Setty et al., 2019].

Pseudotime estimation addresses the task of ordering observed cells into a sequence of cell states traversed by a differentiation process. Typically, the estimated pseudotime values are interpreted as temporal ordering, not capturing the pace of differentiation. It has been suggested that RNA velocity-based pseudotime has the potential to overcome this limitation [Bergen et al., 2020]. While pseudotime estimation might constitute sufficient characterisation of a linear differentiation process, the description of complex processes with more involved topologies such as bifurcations requires an additional step of trajectory inference. Trajectory inference methods seek to infer a representative sequence of states that characterises the possibly multiple differentiation processes in branching or convergent differentiation [Street et al., 2018, Trapnell et al., 2014, Weng et al., 2021, Zhang and Zhang, 2021].

Typical trajectory inference methods are guided by the assumption that phenotypic similarity reflects temporal proximity. However, static expression profiles are ambiguous with respect to the directionality of potential cell state transitions. This ambiguity constitutes a major limitation of pseudotime ordering and trajectory inference, and specifically precludes data driven assignment of root and terminal states without previous knowledge about the process, as well as resolving complex, i.e. cyclical [Mahdessian et al., 2021] or convergent [Burns et al., 2015] process topologies. It has now become possible to estimate transcriptional activity from scRNAseq data via RNA velocity analysis [Manno et al., 2018], consequently enabling the inference of likely transitions between different cell states in a data-driven fashion, ultimately opening the possibility to mitigate the limitations of afore-mentioned reconstruction approaches.

In this work we present Cytopath, a simulation-based trajectory inference approach that takes advantage of RNA velocity. We demonstrate that Cytopath infers accurate and robust cell state trajectories of known differentiation processes with linear, circular, bifurcated, tree-like, and convergent topologies from scRNAseq datasets. We show that Cytopath has the potential to model interlaced processes with different topologies as well as detect regions of transcriptional programme switching. We also assess the ability of pseudotime estimated by Cytopath to represent the biological process time, also referred to as the *internal clock* of a cell. Trajectory inference with Cytopath is superior to and addresses the limitations of both, state-of-the-art trajectory inference approaches as well as recently developed RNA velocity-based methods [Weng et al., 2021, Zhang and Zhang, 2021].

## 2 Results

In this section we present an overview of Cytopath and its trajectory inference performance assessed on six scRNAseq datasets consisting of cellular differentiation processes with various topologies [Manno et al., 2018, Mahdessian et al., 2021, Bastidas-Ponce et al., 2019, Burns et al., 2015, Qiu et al., 2020, Cerletti et al., 2020]. We show that Cytopath successfully recovers known differentiation processes in all datasets with high accuracy. Furthermore, we compare the performance of Cytopath to the best trajectory inference models for each topology [Saelens et al., 2019], which are Slingshot [Street et al., 2018] for tree-like topology, Angle [Saelens et al., 2019] for cell cycle, PAGA (directionality enabled by velocity pseudotime) [Wolf et al., 2019, Bergen et al., 2020] for graph models. We also include comparison to Monocle3 [Cao et al., 2019] as well as two approaches accounting for RNA velocity information, VeTra [Weng et al., 2021] and Cellpath [Zhang and Zhang, 2021]. We demonstrate that the shortcomings of these tools are successfully addressed by Cytopath and that our RNA velocity-based approach enables the resolution of so far intractable topologies.

### Simulation-based trajectory inference with Cytopath

Trajectory inference with Cytopath is performed downstream of the RNA velocity analysis of a scRNAseq dataset, and is specifically based on the resulting cell-to-cell transition probability matrix. The transition probability matrix considers each cell to be a node in a graph and each node is assumed to represent a possible state of the differentiation process under study. The entries of this matrix are the probabilities of transitioning from the a given state to any other state represented in the graph [Manno et al., 2018] [Bergen et al., 2020]. While we base our analysis on an RNA velocity analysis, in principle any cell-to-cell transition probability matrix can be used as input for trajectory inference [Figure 1A.2].

**Figure 1:**
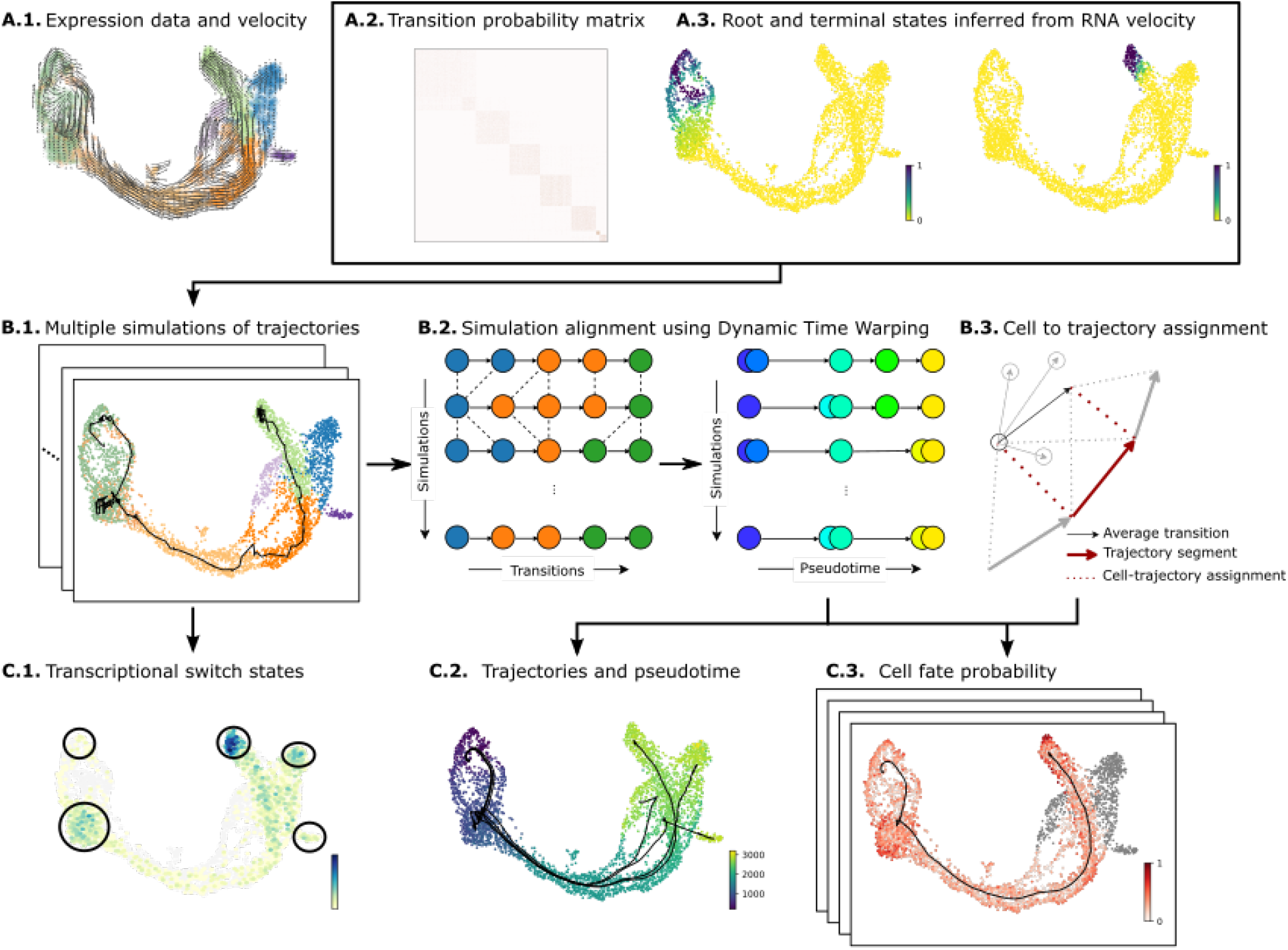
Cytopath overview. **(A)** Inputs for Cytopath trajectory inference subsequent to a RNA velocity analysis. (A.1.) Single-cell gene expression profiles, RNA velocity profiles, (A.2.) Transition probability matrix, (A.3.) Root and terminal state annotation. (shown here: inferred using RNA velocity) **(B)** Steps performed during Cytopath inference. (B.1.) Simulations of the differentiation process generated by sampling a Markov chain based on the cell to cell transition probabilities. Sampling is initialized on cells annotated as root states. (B.2.) Simulations are performed for a fixed number of steps that are automatically selected using the properties of the scRNAseq dataset. Transition steps are aligned using Dynamic Time warping. After alignment cells at each transition step represent the same consensus state. (B.3.) Cells along the inferred trajectory are assigned to multiple trajectory segments based on the alignment of their average transition vector (with respect to neighbours) and the trajectory segment. **(C)** Outputs from Cytopath trajectory inference. (C.1.) The frequency of simulations terminating at each cell highlights regions of switch in transcriptional programs as well as terminal regions. (C.2.) Trajectories are inferred independently for each terminal region. The trajectories are composed of multiple segments. The pseudotime of a cell is estimated as the weighted average segment rank of all the segments it aligns with. (C.3.) Differential alignment scores to multiple trajectories are used to estimate the cell fate probability with respect to the terminal regions.

The objective of trajectory inference with Cytopath is to estimate trajectories from root to terminal cell states, which correspond to the origin and terminus of the differentiation process under study. Root and terminal states can either be derived from a Markov random-walk model utilizing the transition probability matrix itself [Figure 1A.3], as described in [Manno et al., 2018], or can be supplied by the user based on suitable prior knowledge.

The trajectory inference process is divided into four steps [Figure 1B and Methods]. In the first step, Markov sampling of consecutive cell state transitions is performed based on the probabilities of the transition matrix, resulting in an ensemble of simulated cell state sequences. Sampling is initialized at predefined root states and performed for a fixed number of steps until a sufficient number (auto-selected with default settings) of unique cell state sequences terminating within clusters containing the terminal states have been generated [Figure 1B.1]. A pre-computed clustering can be provided to Cytopath to determine terminal regions, otherwise, a clustering is computed internally using Louvain.

The generated cell state sequences are individual simulations of the differentiation process from root to terminal state. Due to the stochastic nature of the sampling process, the cell state sequences cannot be considered as aligned with respect to the cell states at each transition step. Consequently, in the second step, simulations that terminate at a common terminal state are aligned using Dynamic Time Warping which is an algorithm for comparing and aligning temporal sequences with a common root and terminus but possibly different rates of progression. The procedure aligns simulations to a common differentiation coordinate such that cell states from any simulation at a particular differentiation coordinate (pseudotime) represent similar cell states [Figure 1B.2].

Third, consensus expression states across the steps of the aligned simulations are estimated, giving rise to the reported trajectory. Cell states at every step of the ensemble of aligned simulations are averaged and the average value is considered as the consensus state of the trajectory at the particular step [Figure 1C.2]. Alternatively, trajectories can be anchored to observed cell states by choosing the closest cell state to the aforementioned average value. Subsequently, the coordinates of the trajectory with respect to the expression space as well as any lower dimensional embeddings such as UMAP or t-SNE are calculated.

In the final step, cells are assigned to each step of the inferred trajectory. Assignment is based on an alignment score that evaluates for each cell both similarity of its static as well as the velocity profile with each trajectory step. For efficiency, this alignment score evaluation is restricted to cells in the neighborhood around each trajectory step. However the user can optionally compute alignment of every cell for every step of every trajectory. The cell level score is used to estimate position within the trajectory, i.e. pseudotime, as well as the relative association of a cell state to possibly multiple branches of a differentiation processes with complex topology, i.e. cell fate [Figure 1C.2-C.3].

### Reconstruction of neuronal differentiation in the developing mouse hippocampus

We assessed the capability to reconstruct developmental processes with multiple branching which is a frequent topology for scRNAseq datasets generated from experiments studying differentiation processes.To this end, we applied Cytopath and baseline methods to the developing mouse hippocampus dataset, which was first used to demonstrate RNA velocity of single-cells. This dataset is composed of 18,140 cells. The dataset comprises five terminal regions and a common root state. The topology of the data is multifurcating with development branches arising directly from the root state, namely astrocytes and oligodendrocyte precursors (OPC), but also as branches from intermediate states, namely neuroblast and Cajal-Retzius (CA) differentiation.

We recreated the analysis outputs including RNA velocity and the transition probability matrix as indicated in the original publication using scripts made available by the authors. RNA velocity was used to estimate the root and terminal state probabilities [Manno et al., 2018].

Spearman correlation of inferred pseudotime with the cell type identities and their ordering reported in the initial study [Figure 2A] were used for trajectory inference performance assessment [Figure 2D-E]. Cytopath was run using default parameters. It internally selects root and terminal states based on the root and terminal state probabilities estimated in the prior velocity analysis. We also supply the known root and terminal states as supervision to Slingshot and Monocle3 (accepts root states only) to get the best performance from these methods. This could not be done for VeTra and Cellpath since these approaches do not allow inclusion of this supervision. We also assessed the performance of PAGA with velocity-pseudotime-based directionality and *scvelo* latent time. The latter two are not trajectory inference methods, with PAGA only generates a coarse graph of cluster connectivities and latent time is only a pseudotime that does not compute lineage association of cells. However, as RNA velocity-based methods, that have a partial overlap with Cytopath’s core functionality, this comparison is of interest to the community.

**Figure 2:**
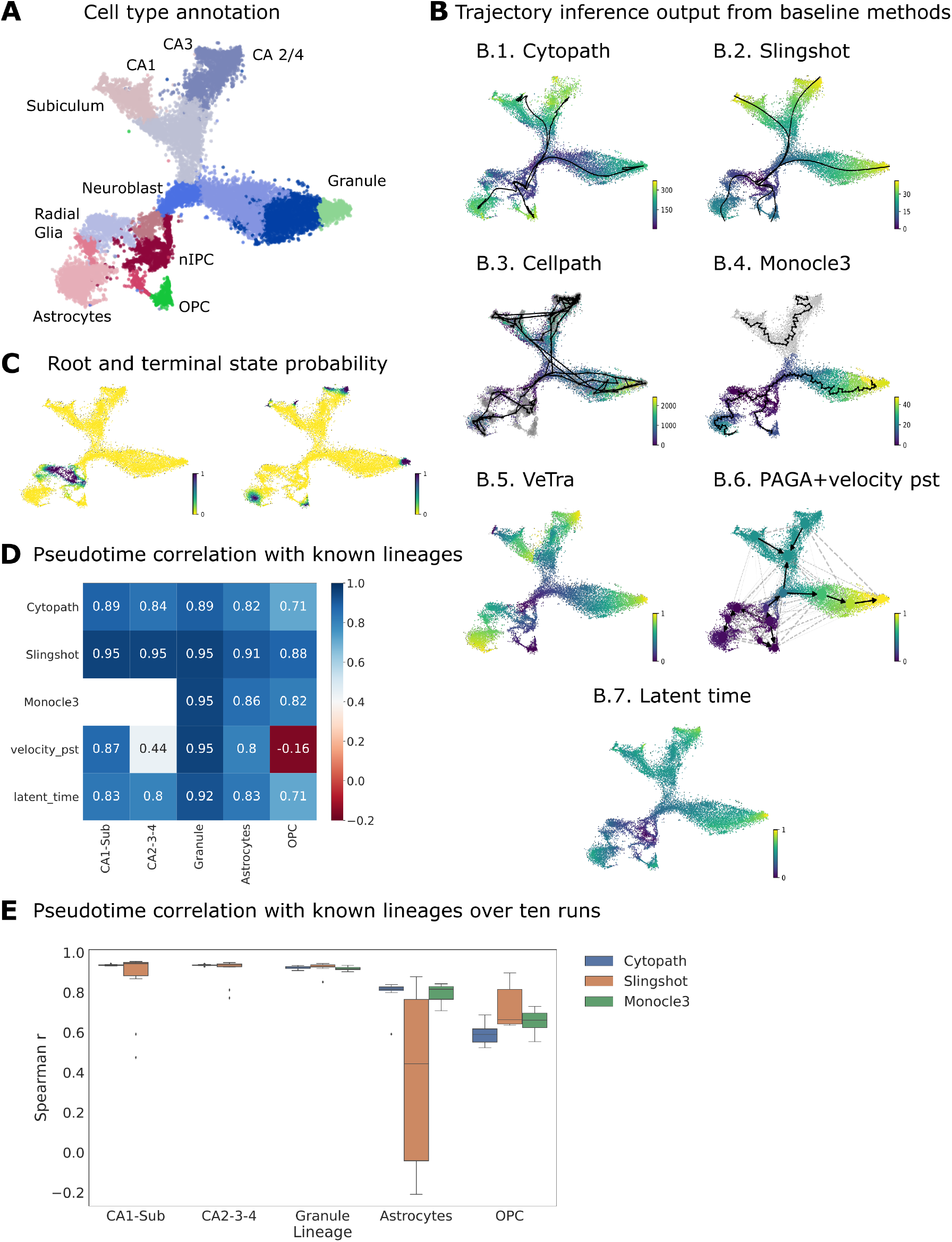
Reconstruction of neuronal differentiation in the developing mouse hippocampus. (A) t-SNE projection of the Dentate gyrus scRNAseq dataset annotated with stages of neuronal differentiation. (B) Trajectory and/or pseudotime inference using (b.1.) Cytopath, (b.2.) Slingshot, (b.3.) Cellpath, (b.4.) Monocle3, (b.5.) VeTra, (b.6.) PAGA+velocity pseudotime (vpt) and (b.7.) *scvelo* latent time. (C) Root and terminal state probability used by Cytopath to select root and terminal regions. (D) Spearman correlation of pseudotime inferred by each method with known ordering of cell types for each lineage. (E) Methods were run ten times to assess the effect of stochasticity in inference (Cytopath) and stochastic estimation of the t-sne embedding (Slingshot and Monocle3)

Trajectories and pseudotime estimated by each method are shown in Figure 2B. Cytopath estimates a single trajectory to each terminal state as expected from known biology. The Spearman correlation between pseudotime inferred by Cytopath for each trajectory to known ordering of cell types is high [Figure 2D] and robust across multiple runs [Figure 2E]. Slingshot also produces trajectories to each terminal state with high correlation however was found to generate one or more spurious trajectories in each run. While the median Spearman correlation of trajectories inferred by Slingshot is high, it appears to have high variability in it’s performance with substantially lower correlation for some runs. This may be due to projection artefacts in the embedding generated in those instances. Monocle3 fails to produce a connected trajectory, producing a disjoint graph and is therefore unable to estimate pseudotime for a large portion of the dataset. Monocle3 pseudotime, velocity pseudotime and latent time are inferred per cell and unlike other methods presented here do not partition cells into trajectories. We used known cell type orderings to select cells relevant to each lineage to perform the correlation analysis.

The velocity-pseudotime method appears to compute a global pseudotime that is incompatible with known differentiation of lineages in this dataset. Consequently, directionality of cluster transitions for PAGA appear to be reversed for CA1Sub and CA2-3-4 lineages and are unclear for OPC and Astrocyte lineages. The latent time method appears to have a similar correlation profile to known lineages as Cytopath [Figure 2D].

Both VeTra and Cellpath infer erroneous trajectories that initiate at terminal or intermediate states. Cellpath generates a large number of trajectories far exceeding the number of known lineages. This is a pattern that is consistent across several datasets and thus a quantitative comparison as performed for other methods presented here is not feasible [Figure S1]. Both methods also exclude a large number of cells from the trajectory inference process. Since VeTra and Cellpath initialise trajectories in intermediate states, cell assignment by these methods does not correspond to a pattern expected for a hierarchical branching structure [Figure S2A.3-A.4].

### Cell cycle reconstruction

We hypothesized that the ability to infer repeating patterns during differentiation likely differentiates RNA velocity- based trajectory inference from other methods that are based on similarity of expression. First, the revisiting the root state implies that inferring the overall direction of the trajectory is not trivial. Second, cells at the origin are a mix of late and early stage state that are co-located in expression space but can be expected to have differing velocities. To assess this hypothesis, we compared the reconstruction of the cell cycle in a dataset comprising 1067 U2OS cells generated using the SMART-Seq2 protocol [Mahdessian et al., 2021].

Based on the comparatively low expression of cell cycle marker genes *Ccne2, Cdk1, Ccna2, Birc5* [Figure 3A-B] we annotated a portion of G0 stage cells (cluster 5) as G1-checkpoint [Figure S4A]. The cell cycle phase annotation per cell from [Mahdessian et al., 2021] was determined using fluorescence intensity of GFP-tagged GMNN (530 nm) and RFP-tagged CDT1 (585 nm). Therefore, the association between cell cycle phase and expression levels of markers is not in phase unlike computational cell cycle phase prediction [Figure S4B-C]. We use phase annotations only to validate the trajectory reconstruction but not for inference. Root and terminal states were selected based on probabilities estimated using *scvelo* [Bergen et al., 2020].

**Figure 3:**
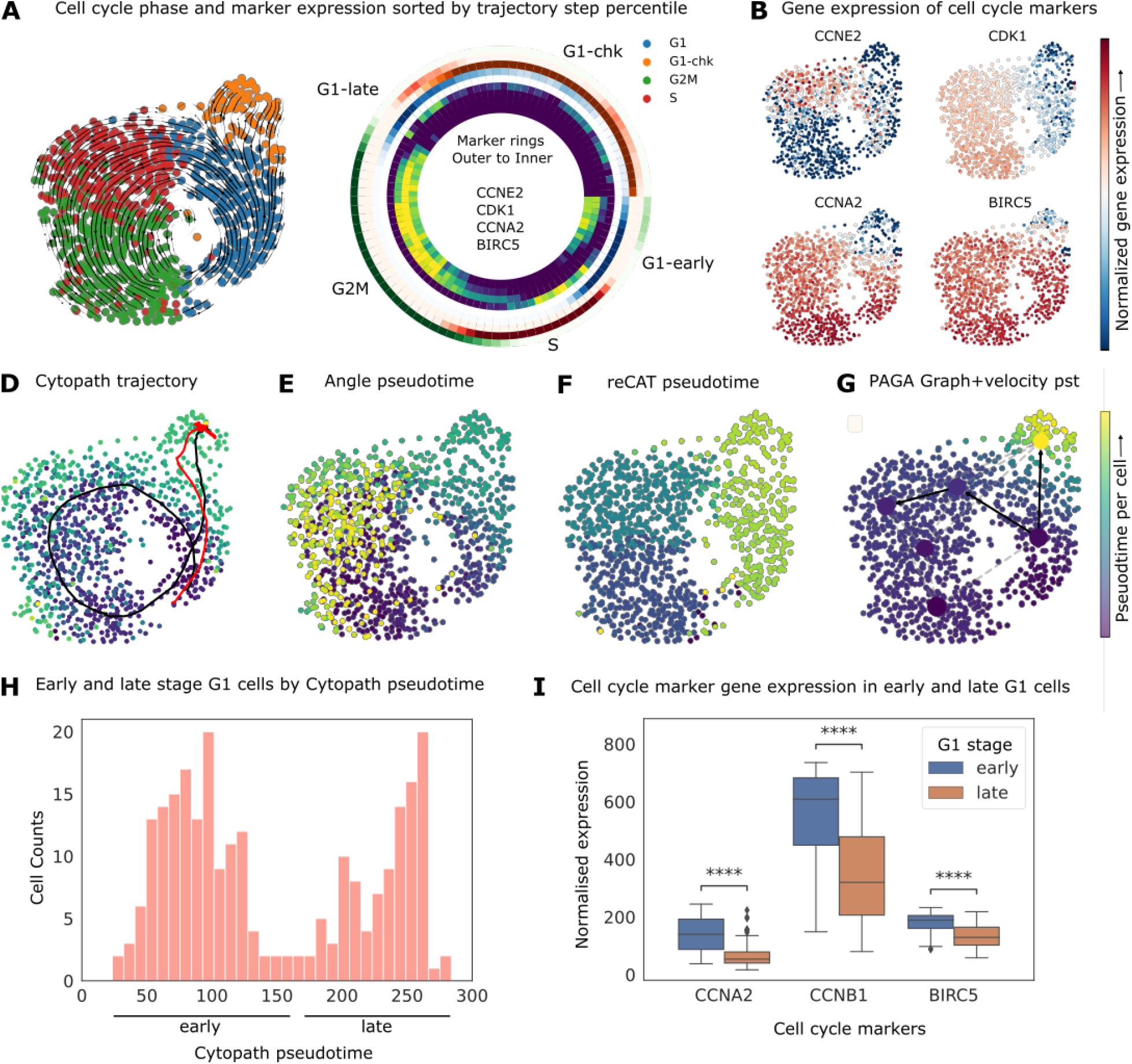
Reconstruction of cell cycle in U2OS cell line. (A) RNA velocity stream plot overlayed on the UMAP projection, annotated with cell cycle phase adapted from [Mahdessian et al., 2021]. Considering all cell-to-trajectory alignments binned into percentiles, the radial heatmap shows cell cycle phase fraction (outer set of rings) and marker expression (inner set of rings) sorted by trajectory step. The directionality of the radial heatmap is clockwise with the origin at zero degrees. The separation of G1 phase into G1 and G1-chk was performed on the basis of (B) marker expression of cell cycle genes. (D-G) Trajectories inferred and pseudotime per cell by (D) Cytopath, (E) Angle, ReCAT, (G) PAGA & velocity pseudotime. (H) Distribution of Cytopath pseudotime for cells in the G1 cluster. (I) Normalised expression of cells classified as early and late G1 cells (blue/orange respectively). Significance was estimated by an independent t-test for each marker.

Cytopath generates a full circular trajectory without interruptions from the cells in G1 stage through the intermediate stages back to the G1 stage and further into the cells in the G1-checkpoint stage [Figure 3D]. The lower expression of relevant cell cycle markers in the terminal region of the trajectory inferred by Cytopath indicates that the pseudotime inferred by Cytopath is a valid representation of the temporal process. Cytopath infers a second linear trajectory indicated in red [Figure 3D]. This is not unexpected as apart from the circular route, a direct connection of the root to the terminal region is also a possible outcome.

Since Slingshot and Monocle3 both assume a tree-like topology they are inherently unsuited to model cyclical trajectories. To compare Cytopath against non-velocity-based baseline methods we selected Angle and ReCAT, both methods intended to model cell cycle in scRNAseq data. In contrast to trajectories inferred by Cytopath, the pseudotime inferred by Angle and ReCAT are inconsistent with marker expression and cell cycle phase annotation. We observe two distinct modes of failure for either method. While pseudotime inferred by Angle represents a partially correct sequence of clusters, the G1-checkpoint cluster is not distinguished from the G1 cluster. Furthermore, S-phase is incorrectly identified as the terminal state. ReCAT successfully identifies G1-checkpoint as the terminal state but detects an incorrect sequence of cell cycle phases [Figure 3E-F].

With respect to the second part of our hypothesis, we observe that cells in the G1 cluster can be divided into two groups on the basis of pseudotime inferred by Cytopath [Figure3H]. Expression of markers associated with cell cycle are significantly higher in the early pseudotime G1 cells than for those destined to move into G1-checkpoint phase and accordingly associated with higher pseudotime [Figure 3I]. Post trajectory inference, cells are assigned to trajectories using the alignment procedure shown in Figure 1B.3. We sort these cell-to-trajectory alignments by trajectory step percentile, then compute cell cycle phase frequency and average marker expression. The partitioning of G1 cells into early and late stage as well as the difference in marker expression can be clearly observed as the two separate bands of G1 (blue) in the radial plot [Figure 3A].

We assessed PAGA with directionality inferred using velocity pseudotime 3G]. PAGA failed to estimate an unbroken sequence of cluster transitions with a default threshold (connections in black) and if the entire connectivity graph is considered (all connections), there are several spurious connections. While the underlying velocity pseudotime is positively correlated with cell cycle phase, G1 cells are not partitioned into early and late stage states. Latent time also models the sequence of cell cycle phases correctly and identifies G1-checkpoint phase as the terminus. However, similar to velocity pseudotime, G1 phase cells are not partitioned into early and late stage states [Figure S3B.3].

Finally, both VeTra and Cellpath which are RNA velocity enabled trajectory inference methods fail to reconstruct the cell cycle correctly. The pseudotime inferred by VeTra appears to be inconsistent with the root and terminal state probabilities. Both methods infer erroneous trajectories that do not capture the cyclical process, both originating and terminating in intermediate states. Both methods do not assign large number of cells to any trajectory [Figures S1B.3-B.4 and S2B.3-B.4].

### Reconstruction of interlaced cell cycling and bifurcated differentiation in pancreatic endocrinogenesis

We further assessed trajectory inference performance for processes with multiple interlaced non-trivial topologies. To this end we considered a dataset studying pancreatic endocrinogenesis with lineages to four terminal states (alpha, beta, gamma and delta cells) and dominant cell cycling at the onset of differentiation [Bastidas-Ponce et al., 2019, Bergen et al., 2020]. Preprocessing, RNA velocity and transition probability matrix estimation were performed with *scvelo* [Bergen et al., 2020] using parameters indicated in the notebook associated with this dataset.

Cell type annotation from [Bastidas-Ponce et al., 2019, Bergen et al., 2020] was used to provide terminal state supervision to Cytopath [Figure 4A] while root states were inferred using RNA velocity. The inferred terminal state probability only identifies the Beta terminal state [Figure S5C]. If the other terminal states are not manually specified then this exclusively data driven approach would report only the trajectory corresponding to the Beta lineage. However, Cytopath can be used to generate undirected simulations not constrained to terminate at a fixed terminal region [Methods]. For each cell, the frequency of simulations terminating at that state can be used to discover regions of transcriptional state switching. Using this approach, two more terminal states (Alpha and Delta) could be recovered [Figure 4C]. However, the trajectory to the Epsilon terminal state could only be constructed by explicitly providing it as terminal state. The set of trajectories estimated by Cytopath corresponding to the four terminal cell types capture the expected differentiation events of endocrinogenesis [Figure 4B].

**Figure 4:**
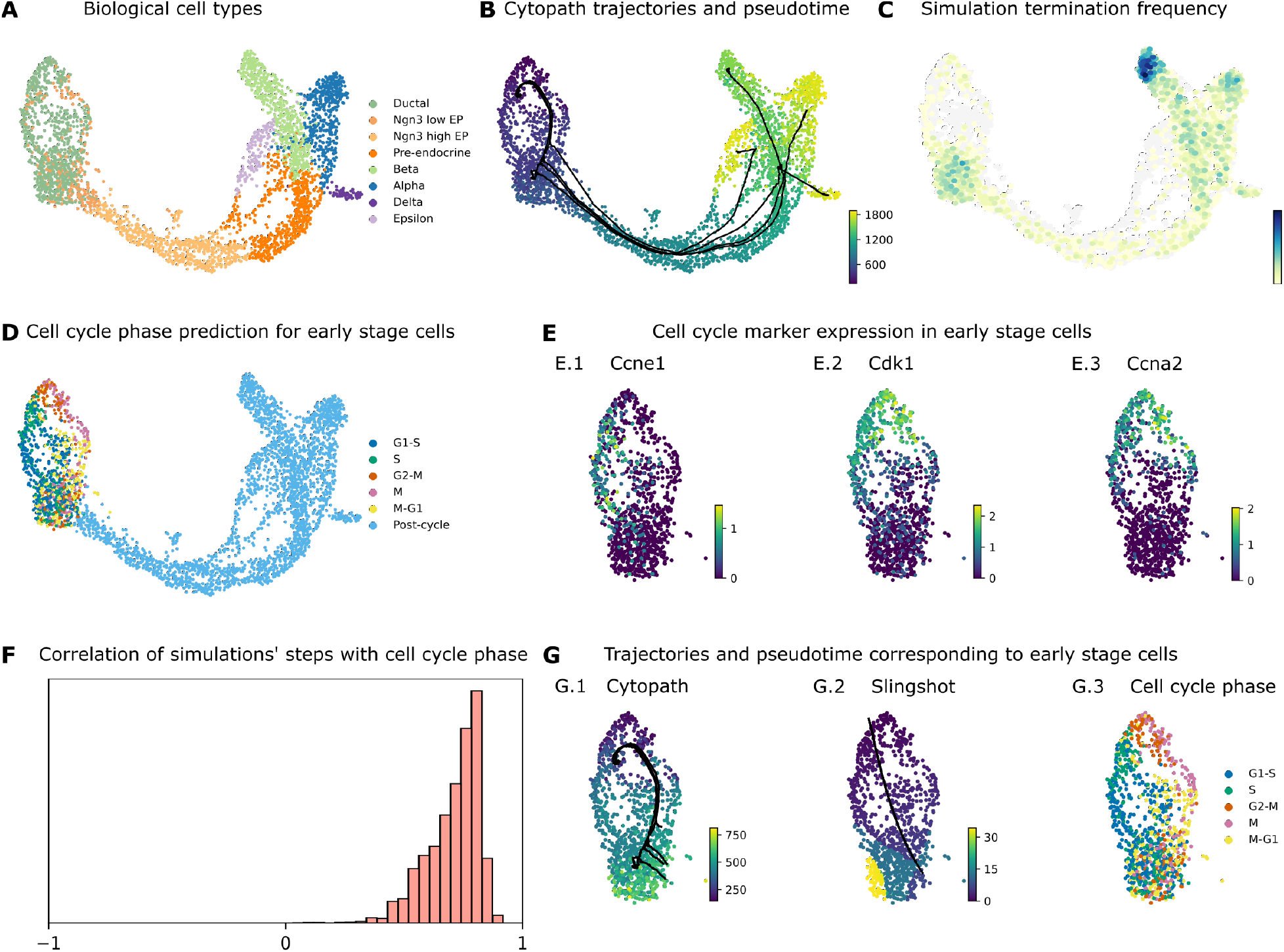
Reconstruction of interlaced cell cycling and bifurcated differentiation in pancreatic endocrinogenesis. (A) UMAP projection of pancreas scRNAseq data annotated with stages of differentiation. (B) Trajectories inferred by Cytopath and mean pseudotime per cell. (C) Log terminal state frequency per cell of undirected simulations initialised at randomly chosen cells. (D) Computational cell cycle phase prediction. (E) Cell cycle marker genes expression in early stage (Louvain clusters 9, 0 and 4) [Figure S6A] (F) Spearman correlation of cell cycle phase with transition step of individual Cytopath simulations. (G) Trajectories and pseudotime inferred by (G.1) Cytopath and (G.2) Slingshot in the root region and (G.3) cell cycle phase annotation.

The trajectories visualised on the UMAP projection indicated a non-linear structure in early stage (root region) cells. Undirected simulations performed using Cytopath also indicate a region of transcriptional switching in Louvain cluster 0 [Figure S6A]. This observation suggests that this differentiation process might structured in two stages, a cycling and a commitment stage. The following inquiries aim at identifying further evidence for this hypothesis.

Cell cycle scoring of cells in the root region was performed and clearly revealed distinct cell cycle states [Figure 4D]. This interpretation is also supported by the differential expression of cell cycle marker genes in the root region [Figure 4E]. The trajectory inferred by Cytopath from the ensemble of 8000 simulations appears to recapitulate the circular structure of the cell cycle [Figure 4G.1]. Spearman correlation of cell cycle phase with the transition steps of each simulation indicate faithful recapitulation of the cell cycle stages at the single-simulation level [Figure 4F].

The simulation-based approach of Cytopath ensures that even in the absence of explicit supervision, cyclic transcriptional patterns are faithfully reconstructed. In contrast, possibly due to the absence of RNA velocity information, the designated root states appear to be isotropic for conventional trajectory inference approaches like Slingshot and therefore they are unable to capture structured transcriptional heterogeneity within this region [Figure 4G.2].

We also show the trajectory estimation with respect to the full pancreatic endocrinogenesis process [Figure S6B-C]. Slingshot and Monocle3 produce spurious or to few trajectories respectively, also when provided with all root and end points. VeTra reports a spurious trajectory that terminates at the Ductal stage, while trajectories to Beta and Alpha are initialised in intermediate or terminal cell states. Both VeTra and Cellpath exclude a large number of cells from the trajectory inference process [Figures S1C, S2C and S3C].

### Reconstruction of convergent differentiation in developing neonatal mouse inner ear

Burns et. al. have shown that the development of hair cells (HC) within the sensory epithelium of the urticle originates from transitional epithelial cells (TEC) via support cells (SC). This study also demonstrated a secondary differentiation path from TEC to HC and put forward the existence of a transitional zone where cells can easily switch fate resulting in two convergent differentiation trajectories [Burns et al., 2015]. This dataset presents a challenge for typical trajectory inference methods given its relatively small size of 157 cells as well as a convergent differentiation topology that violates topology related assumptions of several methods.

Root and terminal state probability estimation using RNA velocity was used to select root and end points. PCA projection of the data was generated as indicated in the original study. Cytopath successfully models the two differentiation trajectories demonstrated in the study. The correlation between known cell type ordering and pseudotime inferred by Cytopath is robust for either lineage [Figure 5D-E].

**Figure 5:**
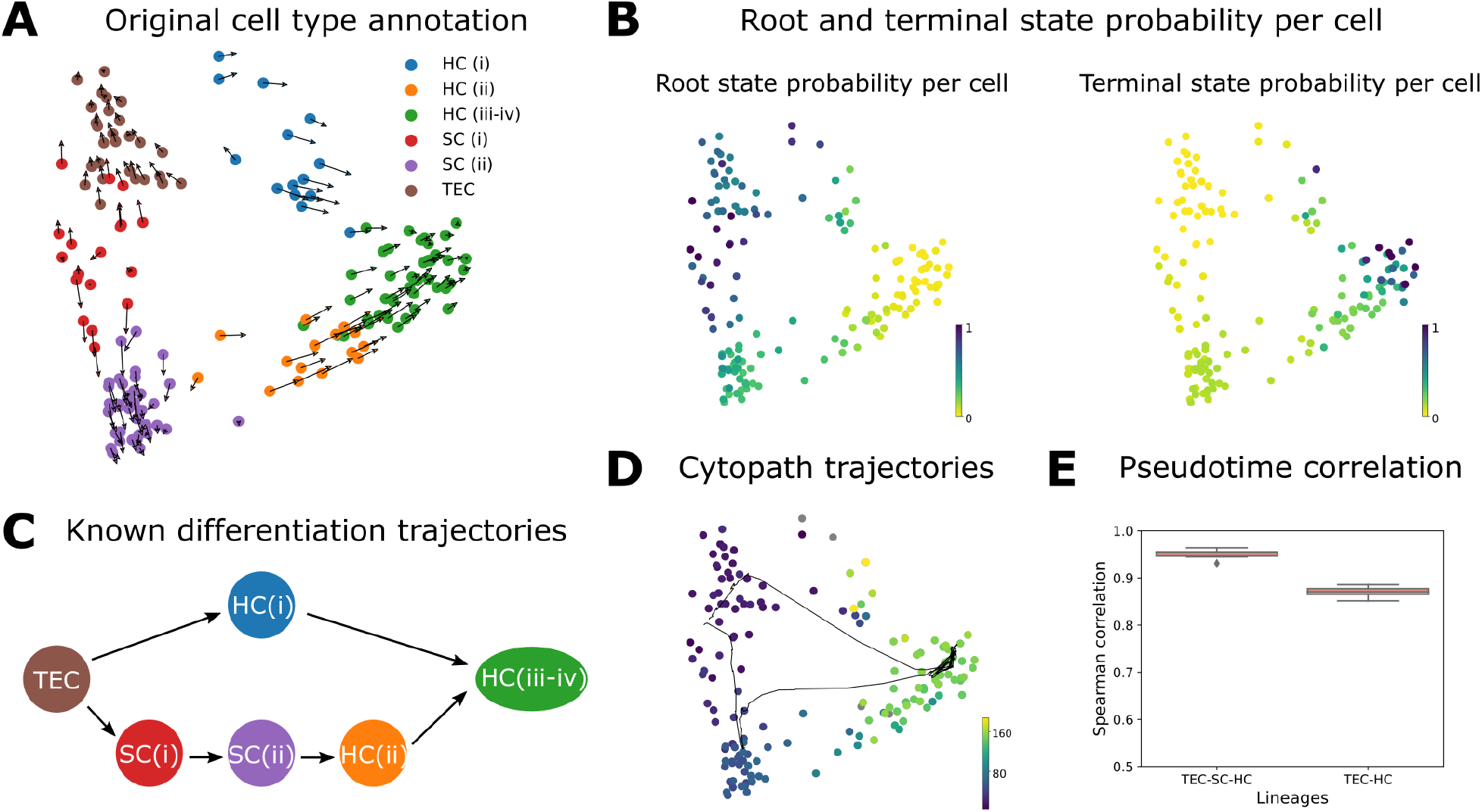
Reconstruction of convergent differentiation in developing neonatal mouse inner ear. (A) RNA velocity overlayed on the PCA projection of neonatal mouse inner ear data annotated with stages of differentiation. (B) Probability estimated based on RNA velocity of a cell being a root state and terminal state respectively. (C) Known differentiation trajectories from [Burns et al., 2015]. (D) Inferred trajectories and mean pseudotime by Cytopath. (E) Spearman correlations between known lineage ordering of cell types and pseudotime inferred by Cytopath (10 runs).

Slingshot, VeTra and Cellpath generate spurious trajectories that terminate at intermediate states [Figure S1D.2-D.4]. None of these methods infers the convergent process. PAGA does not return an unbroken chain of cluster transitions with the default threshold [Figure S3D.4] Monocle3 requires a UMAP embedding and therefore it was not benchmarked in this analysis.

### Cytopath pseudotime inference approximates the internal clock of cells

Difference between two expression states is sufficient to order the cells with respect to progression (difference in expression profiles), however without information on the rate of change of gene expression at any state, the pace of differentiation, i.e. the difference in expression profile relative to the internal clock cannot be inferred. RNA velocity-based trajectory inference and pseudotime inference has the potential to resolve this drawback since RNA velocity provides an approximation of the rate of change of gene expression for each cell.

Single-cell metabolically labeled new RNA tagging sequencing (scNT-Seq) was developed as a means to experimentally measure the age of cells undergoing active transcription. To validate their method, Qiu et. al. generated a dataset of mouse cortical neurons stimulated for durations ranging from 0-120 minutes. The authors also identified a set of activity-regulated genes (ARGs) whose expression can directly be linked to the duration of stimulation. Unlike typical time series scRNAseq datasets where the asynchronous expression of cells implies that the experimental time is decoupled from the internal clock, in the setting described above, the duration of stimulation is a representation of the biological process (internal clock) time with respect to the ARGs. We performed RNA velocity analysis followed by trajectory inference considering only ARG expression. Pseudotime inferred by Cytopath has a monotonic relationship with stimulation time and high Pearson (linear) correlation. To assess the specific relevance of RNA velocity in inferring a pseudotime that better approximates the internal clock, we computed a non-velocity-based pseudotime using the trajectory previously inferred by Cytopath (Cytopath-Euclidean pseudotime). This non-velocity pseudotime has lower median correlation over ten runs than Cytopath pseudotime. Other velocity-based pseudotime estimates also have relatively higher correlation compared to non-velocity-based methods [Figure 6D-E].

**Figure 6:**
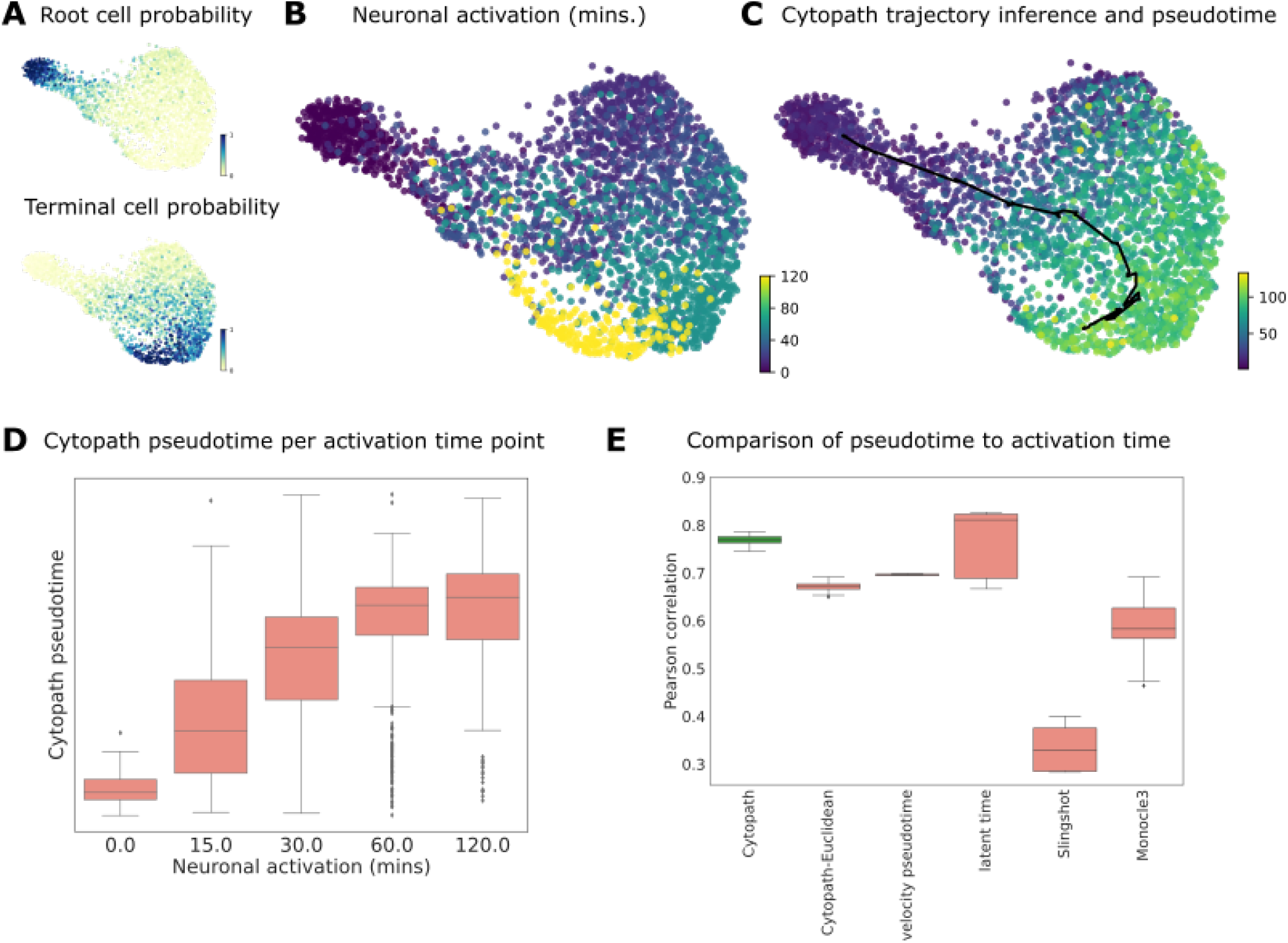
Reconstruction of convergent differentiation in developing neonatal mouse inner ear. (A) Root and terminal state probability inferred using RNA velocity. (B) UMAP projection annotated with duration of stimulation that each cell. (C) UMAP projection annotated with trajectory and pseudotime inferred by Cytopath. (D) Cytopath pseudotime per cell with respect to stimulation duration. Note the monotonic relationship between median pseudotime and stimulation duration. (E) Pearson correlation between pseudotime inferred by Cytopath, non-velocity-based pseudotime inferred using Cytopath trajectory inference (Cytopath-Euclidean) and baseline methods.

In response to stimulation, neuronal cells undergo a relatively fast polarization phase and subsequently slowly return to a depolarized state which is similar to the root state in terms of expression but not rate of change of expression, i.e. RNA velocity. Cytopath-Euclidean pseudotime does not have a monotonic relationship with stimulation duration and places the 120 minute group at a lower pseudotime. We observed the same pattern with Slingshot and Monocle3. However this may partly be due to both poor trajectory inference as well as non-velocity-based pseudotime inference. Surprisingly, latent time and velocity pseudotime which are also RNA velocity-based pseudotime methods also showed a similar pattern of lower pseudotime associated with the 120 minute group of cells [Figure S7].

In the absence of an experimental measure of process time, it is difficult to conclusively explore the association of pseudotime with process time in other datasets presented in this paper. However, if we assume that RNA velocity is a good approximation of transcriptional rate of change, we show that pseudotime inferred using Cytopath outperforms non-velocity-based methods at approximating real rate of change of transcription. The similarity of a cell’s velocity to each of its neighbours indicates the pace of coherent change of transcription in a region of transcriptional space. To quantify this property we define *velocity cohesiveness* [Methods]. High velocity cohesiveness indicates that the cell is present in a region of coherent, and therefore, rapid transcriptional change as the cell has high transition probability to similarly oriented transition partners. Conversely, low velocity cohesiveness indicates that rate of transcriptional change is low and the cell transitions to its neighbours are less coherently directed. Since simulations generated by Cytopath are based on the aforementioned transitions, we expect that the pseudotime estimated by Cytopath better reflects the rate of real transcriptional change compared to a non RNA velocity-based pseudotime that by design is forced to assume a uniform rate of transcription.

We tested this hypothesis by comparing pseudotime estimated using Slingshot and Cytopath for the pancreatic endocrinogenesis dataset. For each lineage we estimated the relative rate of change of Slingshot pseudotime with respect to Cytopath pseudotime per cell. The high positive correlation between velocity cohesiveness and velocity magnitude indicates that in regions with lower velocity magnitude, Slingshot has a lower rate of change in pseudotime as compared to Cytopath and vice versa [Figure S8A-B]. We further define the simulation step density of a cell as the number of unique simulation steps visiting this cell. The negative correlation between simulation density per cell and velocity cohesiveness indicates an enrichment of transitions within regions of slower transcriptional change and vice versa [Figure S8C]. The overall trajectory inferred from these simulations assigns a larger range of pseudotime values to regions with lower velocity cohesion since smaller changes in expression are associated with relatively larger passage of time compared to regions of higher velocity cohesiveness.

### Reconstruction of bifurcating differentiation of CD8+ T cells from scRNA seq time series data

We assessed the performance of Cytopath on a scRNAseq time-series dataset from CD8 T cell differentiation in chronic LCMV infection [Cerletti et al., 2020]. In this infection model system CD8 T cells differentiate from early activated cells into *exhausted* and *memory-like cells* over a period of three weeks. Samples were collected at eight experimental time points after infection with LCMV to cover all stages of the process [Figure 7A]. Although these samples are heterogeneous snapshots of a spectrum of differentiation states at a particular time point, they provide an approximate development coordinate. Starting from a population of cells at an early activated state, differentiation leads into the two distinct terminal states within five days of the LCMV infection. This differentiation process is characterized by strong transcriptional changes and expression of different surface genes.

**Figure 7:**
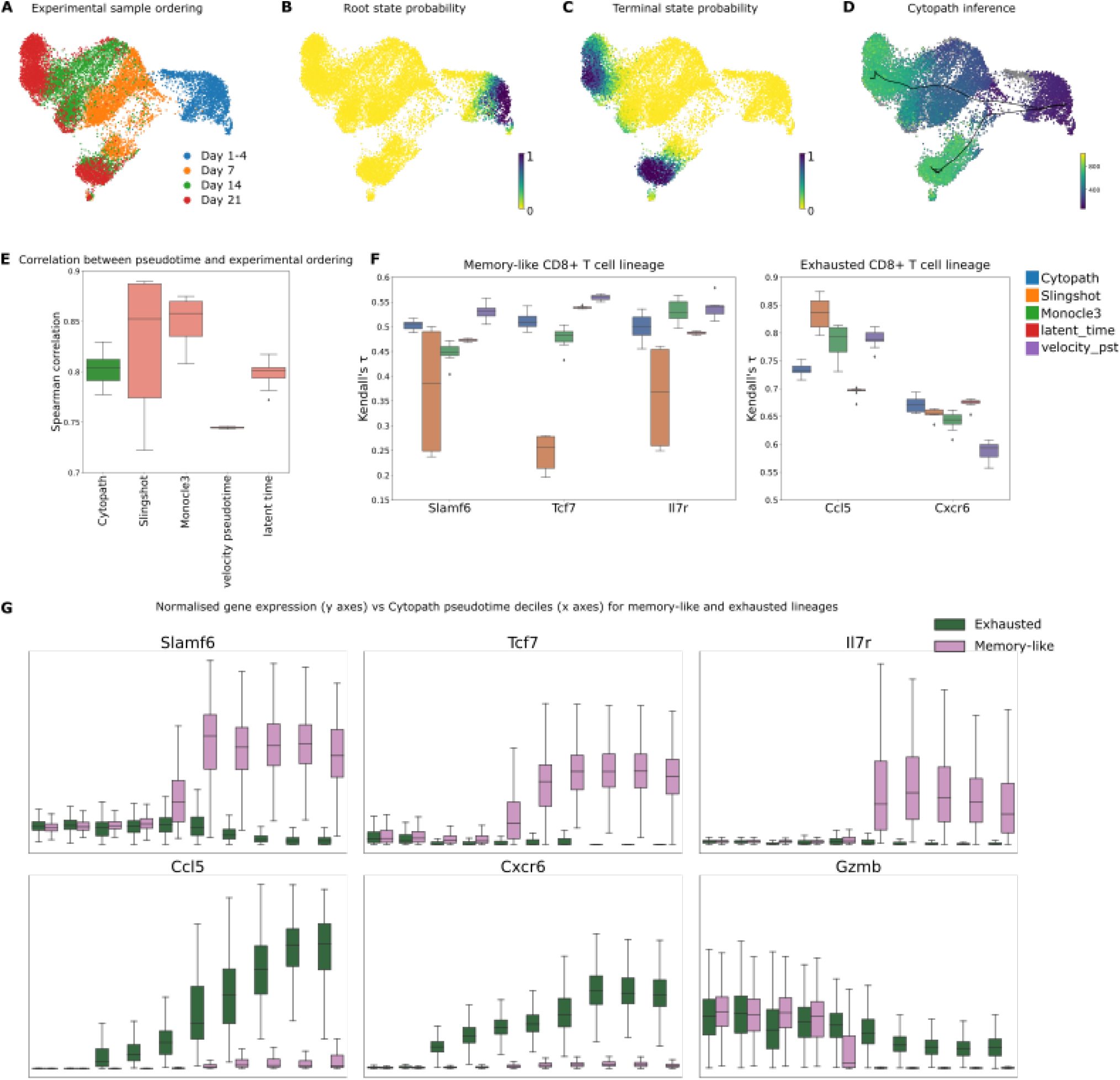
Reconstruction of bifurcating development of CD8+ T cells from scRNA seq time series. (A) Ordering of samples with respect to time of LCMV innoculation. (B-C) Probability estimated based on RNA velocity of the cell being (B) a root state (C) a terminal state (D) Trajectories inferred and mean pseudotime per cell by Cytopath.(E) Correlation of pseudotime estimated by each method with marker relevant to the Memory-like CD8+ T cell differentiation and Exhausted CD8+ T cell differentiation respectively. (F) Normalized expression of key marker genes in each Cytopath pseudotime decile per lineage. First row (*Slamf6, Tcf7, Il7r*) markers are expected to be expressed in the Memory-like lineage. *Ccl5* and *Cxcr6* are expected to be expressed only in the Exhausted lineage. *Gzmb* is not expected to be differentially expressed between the two lineages.

We identified the root and terminal states of the process using *scvelo* [Bergen et al., 2020] [Figure 7B-C]. The terminal states were validated by expression levels of known marker genes [Cerletti et al., 2020]. The *exhausted* terminal state showed high expression in co-inhibitory markers like CD39 (*Entpd1*), CD160 (*Cd160*) and PD-1 (*Pdcd1*). The *memory-like* terminal state had high expression of TCF1 (*Tcf7*) and IL-7R (*Il7r*). Trajectory inference with Cytopath resulted in two trajectories that lead from a shared starting region to the two expected terminal regions. The two trajectories overlap in the beginning of the process but then sharply diverged at a branching point [Figure 7D].

Comparing the pseudotime estimates from Cytopath with the discrete experimental time labels from the samples showed high agreement of the two. The experimental time labels, corresponding roughly to the developmental coordinate, were ordered correctly and with high Spearman correlation between pseudotime and time labels [Figure 7E].. It is important to note that unlike analysis of the neuronal activation dataset’s activity-regulated genes, the experimental time in this setting is not a precise representation of the internal clock of every cell. This is due to asynchronous activation of expression arising from cell-to-cell variation of antigen exposure times. Therefore we only expect a correct ordering of experimental time labels with respect to median pseudotime per experimental sample and not a perfect correlation of pseudotime and experimental time label per cell.

Cytopath suggests a bifurcating trajectory model with trajectories originating at biologically relevant root states and terminating at either of the expected terminal states [Figure 7D]. Slingshot also inferred trajectories to either terminus but generated a third spurious trajectory within the early stage cell group that cannot be matched to any expected infection-induced differentiation trajectory [Figure S1F.2]. VeTra and Cellpath infer multiple erroneous trajectories that do not initiate at the root state and estimate pseudotime that does not correspond to the differentiation process at all [Figure S1F.3-F.4,S2F.3-F.4]. Monocle3 reconstructs the global structure of the data, but includes additional loops and branches within the exhaustion branch [Figure S3F.2].

Further, we assessed the correlation of pseudotime estimates with canonical gene expression markers of the memory-like (*Slamf6, Tcf7, Il7r*) and exhaustion branch (*Ccl5, Cxcr8*). We observe that pseudotime inferred by Cytopath is highly correlated with lineage specific markers [Figure 7F].

We further tested the validity of the average trajectories of Cytopath by the expression profiles of known lineage marker genes in the differentiation process. The chemokine receptor CXCR6 has been shown to mark exhausted T cells in chronic LCMV infection [Sandu et al., 2020]. The average expression of *Cxcr6* increases in the trajectory towards the exhausted cluster just prior to the divergence of the two branches [Figure 7G], indicating that the paths are indeed governed by the exhaustion process. Conversely T cell Factor 1 (TCF1) and the expression of its gene *Tcf7* is an established marker of memory-like cells [Utzschneider et al., 2016]. Expression of this gene was increased in memory-like cells just after the cells started to diverge after the bifurcation point. Towards the memory-like terminal state at late time-points, *Tcf7* expression is exclusive to the memory-like population. An additional observation is the high expression of *Gzmb* early in both branches that drops off towards later timepoints [Figure 7G]. The expression of *Gzmb* is a shared feature of both branches and known to decrease in both branches as the infection progresses and expression is low towards late timepoints [Wherry et al., 2007].

In summary, Cytopath is able to reconstruct biologically relevant differentiation trajectories from a long-term time series dataset, in a more accurate and reproducible manner than widely used tools. We identified correct differentiation branches of CD8 T cells in chronic infection, demonstrated by correct ordering of the experimental time labels and expression levels of branch specific gene expression markers. For this system several phenotypic populations and characteristic markers had been described before, but the connecting differentiation trajectories of those populations are a subject of ongoing research [Chen et al., 2019, Zander et al., 2019, Yao et al., 2019, Raju et al., 2020]. These studies provide evidence for branching in the development process, and only recently in conjunction with simulation-based trajectory inference it was possible to resolve this event in more detail [Cerletti et al., 2020].

## 3 Discussion

Trajectory inference is a challenging task since scRNA seq data is noisy and - until recently - has been evaluated to achieve only static expression profile snapshots. Trajectory inference tools typically operate in low dimensional embeddings, especially two dimensional projections such as UMAP and t-SNE, possibly obfuscating complex trajectory topologies such as multifurcations and cycles due to more dominant sources of variation. Inclusion of directional transcriptional activity estimates from RNA velocity analyses is expected to achieve more precise and sensitive trajectory inference. With Cytopath we present an approach that takes advantage of this information. We demonstrate superior capability to reconstruct complex differentiation hierarchies from diverse scRNAseq datasets with respect to number of cells, differentiation topologies and sequencing technologies.

The transition probability matrix used to generate simulations is computed from high dimensional gene expression and velocity profiles. Since Cytopath is based upon transitions that use the full expression and velocity profiles of cells, it is less prone to projection artifacts distorting expression profile similarity. In addition, this approach specifically considers likely and discards unlikely transitions, and thereby is able to identify for instance cyclic trajectories in an apparently diffusely populated and isotropic region of expression space [Figure 3, 4]. Furthermore, these hidden transcriptional patterns are made apparent by the simulation-based approach without any explicit supervision. Non-RNA velocity-based methods struggle to discriminate between cells corresponding to different stages or branches of cyclical and convergent processes respectively since the cells appear to be co-located in expression space. However, RNA velocity-based pseudotime and cell fate estimation based on analytical analysis of Markov chain properties, performed by *scvelo* and *CellRank* [Lange et al., 2022] also does not readily present this information to the user even if the pseudotemporal ordering or cell fate scoring estimated by these tools captures these patterns.

Cytopath analysis requires specification of the root and terminal regions. This requirement is met easily when the cellular process is sufficiently well characterized up to the level of *a priori* definition of these regions. However, even if this is not the case, Cytopath can detect and utilize tentative root and terminal states from the cell to cell transition probability matrix using *scvelo* [Bergen et al., 2020]. Furthermore, simulations generated using Cytopath can also aid in the identification of terminal cell types. Compared to the absorbing Markov process based inference of terminal states implemented in *velocyto* and *scvelo*, this approach appears to highlight more terminal states in the pancreatic dataset [Figure S9C]. Intermediate quasi-stationary cell states induced for instance by a switch of transcriptional programs appear to be highlighted by this procedure, as indicated for the switch from the cell cycle to islet cell differentiation in the pancreatic endocrinogenesis study. While this simulation-based estimation of root and terminal states is not a core functionality of the presented method, it opens Cytopath’s application to less characterized datasets. In subsequent work, Cytopath simulations could be the basis for learning mechanistic differentiation models and gene regulatory networks.

We also assessed the utility of Cellrank as a means to identify root and terminal states to be used as input for Cytopath trajectory inference and found that - for the datasets considered in this study - Scvelo performed better at recovering known root and terminal states, only failing to identify terminal states in the pancreatic endocrinogenesis dataset. Conversely, Cellrank appears to misidentify root and terminal states in every dataset. [Figure S5, S13 and S14].

While Cytopath is primarily a trajectory inference tool, we leverage the alignment-based association of cells to inferred trajectories to generate additional results that overlap in functionality with other RNA velocity-based tools. For instance, Cytopath can also be used to predict cell fate [Figure S10-S12]. Partition-based graph abstraction (PAGA) [Wolf et al., 2019] is yet another popular tool that can include RNA velocity to infer directed connectivity between clusters of a scRNAseq dataset. Subsequently, PAGA requires a user-defined threshold to separate genuine and spurious connections. Cytopath’s compositional clusters for each trajectory offer an alternative estimate of cluster orderings and can also be used to generate a coarse graph of cluster connectivity. Unlike PAGA, Cytopath does not require a post inference selection of valid connections between clusters that maybe induce confirmation bias. Velocity pseudotime allows directed edges to be inferred using PAGA, however, an unbroken sequence of connections is not guaranteed [Figure S3A.4-F.4]. The coarse graph approach has a few disadvantages compared to trajectory inference methods. Dedicated trajectory inference methods such as Cytopath, Slingshot and Monocle3 can model gradual divergence of lineages. Cell fate scoring estimated by Cytopath constitutes fuzzy assignment of cells to multiple lineages. These methods also support relatively diffuse regions of branching. In contrast PAGA considers Cells within a cluster to be homogeneous with respect to lineage assignment and therefore branching can only be defined at cell cluster level.

Other RNA velocity-based trajectory inference tools, VeTra and Cellpath, appear to perform significantly worse than non-velocity-based methods used as benchmarks in this study [Figure S1-S2]. We assume that the reason for the difficulty to recapitulate trajectories could be their in-built lack to guide trajectory inference by separately providing root and terminal states. This appears to be a contrived problem since biological knowledge regarding the identity and role of cells is typically available, or - as discussed before - can be estimated separately. Ignoring this information seems to make trajectory inference unnecessarily difficult and could be the reason why Cytopath, as well as Slingshot and Monocle3, perform better than VeTra and Cellpath. Regarding trajectory inference with Cytopath, we find that automatic selection of root and terminal states tends to match biologically relevant root and terminal states with the strong exception of the pancreatic endocrinogenesis dataset. In general, we recommend that root and terminal state selection should be done by synthesizing all available sources of information including application relevant cell type marker expression profiles, analytically derived probabilities based on RNA velocity and simulations using Cytopath.

The addition of RNA velocity is expected to allow for pseudotime inference that is a better representation of the *internal clock* of the cell that corresponds to the pace of differentiation. We show that pseudotime inferred by Cytopath has a monotonic relationship with the process time. The three points of evidence we show in this paper are first, the ability to partition cells in the G1 phase of the cell cycling dataset into late and early stage cells. From the perspective of gene expression profiles these cells are co-clustered and appear as a single-cell type. However, the RNA velocity-based cell-to-trajectory alignment procedure implemented in Cytopath assigns these cells to either an early trajectory step corresponding to G1-S phase transition or late stage indicating G1 to G1-checkpoint transition. The biological relevance of this partitioning can be validated by the significant difference in gene expression of a selected set of cell cycle marker genes [Figure 3A]. Second, we investigate in more detail the correlation of pseudotime inferred by Cytopath with stimulation duration for the neuronal activation dataset. By restricting the analysis to ARGs whose expression is triggered in response to stimulation and hence synchronized per cell, we can consider the experimental time ordering to be coupled to the process time in this dataset [Figure 6]. Last, we examine the relationship between velocity magnitude and rate of change of pseudotime. Intuitively, regions with high velocity magnitude are expected to have relatively larger change of expression with respect to the internal clock of cells and vice-versa. We show that this relationship is better modelled by Cytopath than non-velocity-based methods [Figure S8]. These results suggest that pseudotime estimated by Cytopath is an improvement on approximating of real rate of change of gene expression.

We show that pseudotime inferred by Cytopath is robust to root and terminal probability thresholds for root and terminal state selection [Figure S15]. Cytopath’s runtime predominantly scales with the number of simulation steps [Figure S16]. Cytopath considers generic properties of scRNAseq datasets such as total number of cells, number of inferred root and terminal states to initialise the hyper-parameters of the trajectory inference process. This selection is done with the objective of computational efficiency as well as robust detection of trajectories. All the analyses presented in this paper utilized the default automatic hyper-parameter selection approach, however users still retain the option of performing manual tuning.

In summary we expect simulation-based trajectory inference approaches like Cytopath to enable sufficiently precise and unambiguous trajectory inference to achieve testable hypotheses to identify drivers and derive mathematical models of complex differentiation processes.

## 4 Methods

### Trajectory inference with Cytopath

#### Cell clustering

Any grouping of cell states, such as clustering from widely used community detection algorithms or cell type annotations generated by domain experts can be provided as input to Cytopath.

For the set of all cells *C* and the set of all clusters *S*, the clustering is *f*^*s*^ : *C → S* where |*S*| *<* |*C*|.

By default Cytopath will perform clustering of cells using Louvain via *scvelo* [Bergen et al., 2020].

#### Stationary state selection

Root (*P*_*r*_)and terminal (*P*_*t*_) state probabilities are used to determine the stationary states as described previously [Manno et al., 2018, Bergen et al., 2020] By default, a threshold of 0.99 is used.

The set of root cells *C*^*r*^ is defined as *{c ∈ C* : *P*_*r*_(*c*) *≥* 0.99*}* Similarly, the set of all terminal cells *C*^*t*^ is defined as *{c ∈ C* : *P*_*t*_(*c*) *≥* 0.99*}* Terminal regions *S*^*t*^ are defined as *{f*^*s*^(*c*) *∈ S* : *c ∈ C*^*t*^*}*

When prior biological knowledge regarding the data exists, users can also manually specify root and terminal cell states or designate entire clusters as root or terminal regions.

#### Simulations

Simulations are initialized at random cell states selected uniformly within the defined root cells and consist of a fixed number of cell to cell transitions.

At each step, a single transition from the current cell state is realised based on the cell to cell transition probability matrix *P*. Each row of the matrix contains the probability of transition from the current state (row index) to another cell in the dataset (column index). The cell state *c*_*ij*_ at step *i* of simulation *j* is selected randomly according to *P* from the nearest neighbours of *c*_(*i−*1)*j*_.

Let *F* be the cummulative probability distribution of *P*. A value *κ* is sampled from a uniform distribution over [0, 1) and,

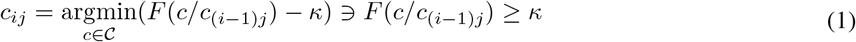

#### Technical parameter selection

Under default settings, the number of simulation steps and minimum number of simulations to be generated are automatically adjusted based on the size of the dataset and number of terminal regions. The purpose of this adjustment is to make the sampling process computationally efficient. The scaling parameters were determined by empirical testing.

The number of simulations steps *i*_*max*_ is initialised as ⌈ 5 log_10_(|*C* |) ⌉. This represents an increase of five simulation steps per order of magnitude increase in the size of the dataset. The minimum number of unqiue simulations to be generated per terminal region *m* is selected as ⌈ 500 *∗* log_10_(|*C*|)⌉

Subsequently the number of simulation steps and number of simulations to be sampled are adjusted during the sampling process in an iterative fashion based on the proportion of simulations that terminate at terminal regions in the previous iteration, until the minimum number of unique simulations per terminal region have been generated.

Let *J* be the set of all simulations generated in an iteration and *J*^*t*^ be the set of simulations terminating in terminal regions. If |*J*^*t*^| *<*= 0.1 *∗* |*J* | then the number of simulation steps is doubled.

Let 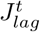 be the set of simulations terminating at the terminal region with least number of simulations, 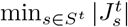. If 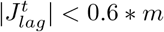 then the number of simulation steps are incremented by 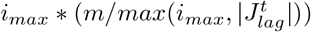

If the two conditions above are met and the minimum number of unique simulations per terminal region are not obtained for any terminal region then more samples are generated until 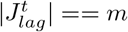 .

#### Trajectory inference

Simulations that terminate within terminal regions are considered for trajectory inference. Trajectory inference is performed by first clustering the simulations and then aligning them using Dynamic Time Warping, which is an algorithm that allows alignment of temporal sequences with a common origin and terminus that possibly have different rates of progression.

For any two simulations *A* = *{c*_0*a*_…*c*_*ia*_*}* and *B* = *{c*_0*b*_…*c*_*ib*_*}*, the Euclidean Hausdorff distance *H*(*A, B*) is defined as,

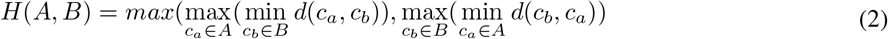

where d is Euclidean distance. Simulations terminating within a single terminal region 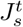 for *s ∈ S*^*t*^, are clustered using Louvain based on Euclidean Hausdorff distance.

Each cluster of simulations is then aligned in a greedy-pairwise fashion using the *fastdtw* python package to generate a single ensemble sequence per cluster which we refer to as a sub-trajectory [Salvador and Chan, 2007].

The mean value of coordinates of cells at each step of the aligned simulations is the coordinate of the (sub-)trajectory at that step. Subsequently, a second round of clustering and alignment is performed, using the sub-trajectories from the first round to produce trajectories that are reported by Cytopath. By default the number of expected trajectories per terminal region is not specified allowing from unbiased inference of multiple trajectories per terminal region. However, users have the option to manually enforce the number of trajectories to infer per terminal region.

#### Identification of compositional clusters

For each trajectory compositional clusters are identified from the set of clusters provided in the step *cell clustering*. For each step *i* of the trajectory, its neighborhood *M*_*i*_ in PCA space is recovered with a K-dimensional tree search [Virtanen et al., 2020]. Cell clusters with a representation larger than a threshold frequency (Default:0.3) for at least one *M*_*i*_ are considered compositional clusters of that trajectory.

#### Alignment score

After the trajectory coordinates have been inferred, the cell-to-trajectory association is computed. Trajectories inferred by Cytopath are segmented and cells in the compositional clusters of a trajectory are aligned to the segments of the trajectory.

For a cell with neighbours *K*, its alignment score to step *i* of a trajectory is the maximum of two scores. The score with respect to the trajectory segment *b* from steps *i-1* to 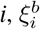 is calculated as,

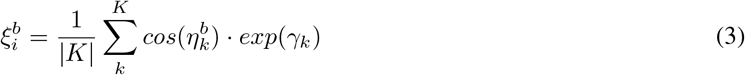

where *η* is the cosine angle between the section of the trajectory and all possible transition partners *k* of the cell. *γ* is the cosine similarity between the velocity vector of the cell with the distance vector between the cell and its neighbours. The score with respect to the trajectory segment *f* from steps *i* to 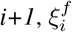 is calculated similarly,

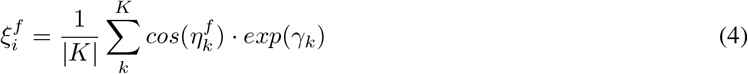

The alignment score is an extension of the transition probability estimation implemented in *scvelo* by weighting each transition of a cell with respect to its alignment to a trajectory. The score is used for assessing the position of a cell with respect to a trajectory (pseudotime) and subsequently to compute a fate score with respect to multiple lineages. The alignment score *p*_*i*_ of the cell with respect to step *i* is 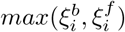

For each cell the final alignment score *p* with respect to a particular trajectory is an average of it’s alignment scores to multiple step segments of the trajectory. By default, it is the mean, however other averages can also be used.

#### Cell fate score

For each cell its alignment score, relative to multiple trajectories is the cell fate score. Cell fate score *f*_*traj*_ for cell with respect to a trajectory is

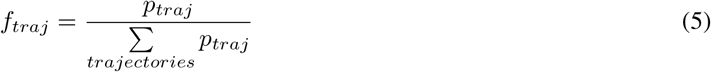

#### Pseudotime estimation

For each trajectory cells that have an alignment score greater than zero and also belong to a compositional cluster of the trajectory are assigned a pseudotime value with respect to the trajectory. Since a cell can align to multiple steps within a trajectory, the mean step value of a cell weighted by alignment score is taken as its pseudotime value, for each trajectory. Optionally, other averages can also be used.

### Evaluating dynamical properties of Cytopath pseudotime

#### Terminal state identification using undirected simulations

For each dataset, five thousand simulations were initialized at randomly chosen cells from the dataset and sampling was performed for 30 steps. The log of total count of simulations, from all the clusters, terminating at each cell, rescaled to range [0,1] is reported as propensity for constituting either a terminal state or an intermediate state representing a switch in transcriptional programme.

#### Cytopath-Euclidean pseudotime

Cytopath-Euclidean pseudotime was computed by considering all the cell-to-trajectory alignments inferred by Cytopath. The cell was assigned to the trajectory segment with minimum Eucldiean distance between the cell and the segment in PCA space.

#### Velocity cohesiveness

Cosine similarity of the distance vector between a cell to its transition partners, and the cell’s velocity vector, per cell is stored as the velocity graph. Velocity cohesiveness of cells are the mean values of each row of this matrix.

#### Pseudotime comparison

Cells were ordered by Cytopath pseudotime. For each cell a window of 50 cells centered on the cell was considered. For each window a linear model was fit with respect to Slingshot and Cytopath pseudotime using *scipy*.*stats*.*linregress* function. Rate of change of Slingshot pseudotime vs Cytopath pseudotime was assessed by estimating the slope.

#### Simulation step density

Simulation step density per cell is the log of the count of simulation steps that are a transition to the cell.

#### Runtime analysis

*process_time* function from the *time* package was used to record the time taken for each trajectory inference procedure.

### Comparison of trajectory inference approaches

#### Parameter settings

Default settings were used for all methods except VeTra and Cellpath. Recommended settings based on information published by the authors were used for VeTra and Cellpath. Cytopath was run using default parameters for all datasets. The same root and terminal states used for trajectory inference with Cytopath were provided to other methods. Figure S5 shows the root and terminal state probabilities estimated using *scvelo* for each dataset.

#### Pseudotime comparison

Spearman correlation of the pseudotime values generated by each method with the cell type cluster ordering for each biological lineage was used to compare the performance of the methods. Kendall’s tau was used to assess the correlation of marker expression with the estimated pseudotime. For each dataset and method, analysed with the correlation analysis described above, the analysis was performed on the dataset with ten independent initialisations of the entire processing pipeline.

### Datasets

Pre-processed data was used wherever available. Subsequent analysis was performed with *scvelo* using standard workflow. Trajectory inference analysis with Cytopath including pre-processing and velocity analysis for each dataset presented in this paper can be found at https://github.com/aron0093/cytopath-notebooks. Download links for *anndata* objects for each dataset are also available in the corresponding notebook.

#### Dentate Gyrus

Prepossessing was performed as indicated by the authors of the original study using code published by La Manno et. al.

#### Neonatal mouse inner ear

Raw sequencing data was downloaded from the NCBI GEO database under accession code GSE71982. Quality control including read filtering and adaptor trimming was performed using *fastp* [Chen et al., 2018]. Reads were aligned to the GRCm39 mouse genome assembly using *STAR version=2*.*7*.*8a-2021-04-2* [Dobin et al., 2012]. Spliced and unspliced counts were estimated using the *velocyto run-smartseq2* command following the recommendation of the developers.

#### CD8 development

Transgenic P14 CD8 T cells were sampled longitudinally during chronic infection with LCMV Clone-13 infection. The samples were acquired from four phases of the infection, namely activation (day 1-4), effector (day 7), early exhaustion (day 14) and late exhaustion (d21) and scRNAseq was performed using the 10x Genomics platform.

Read counts were realigned and sorted for spliced and unspliced counts using the analysis pipeline from velocyto [Manno et al., 2018]. Other contaminating cell types were removed from the dataset based on outliers in diffusion components. [Cerletti et al., 2020]

## Code Availability

Cytopath has been implemented as a python package and can be found at the following GitHub repository (https://github.com/aron0093/cytopath).

## Data Availability

**Table 1:**
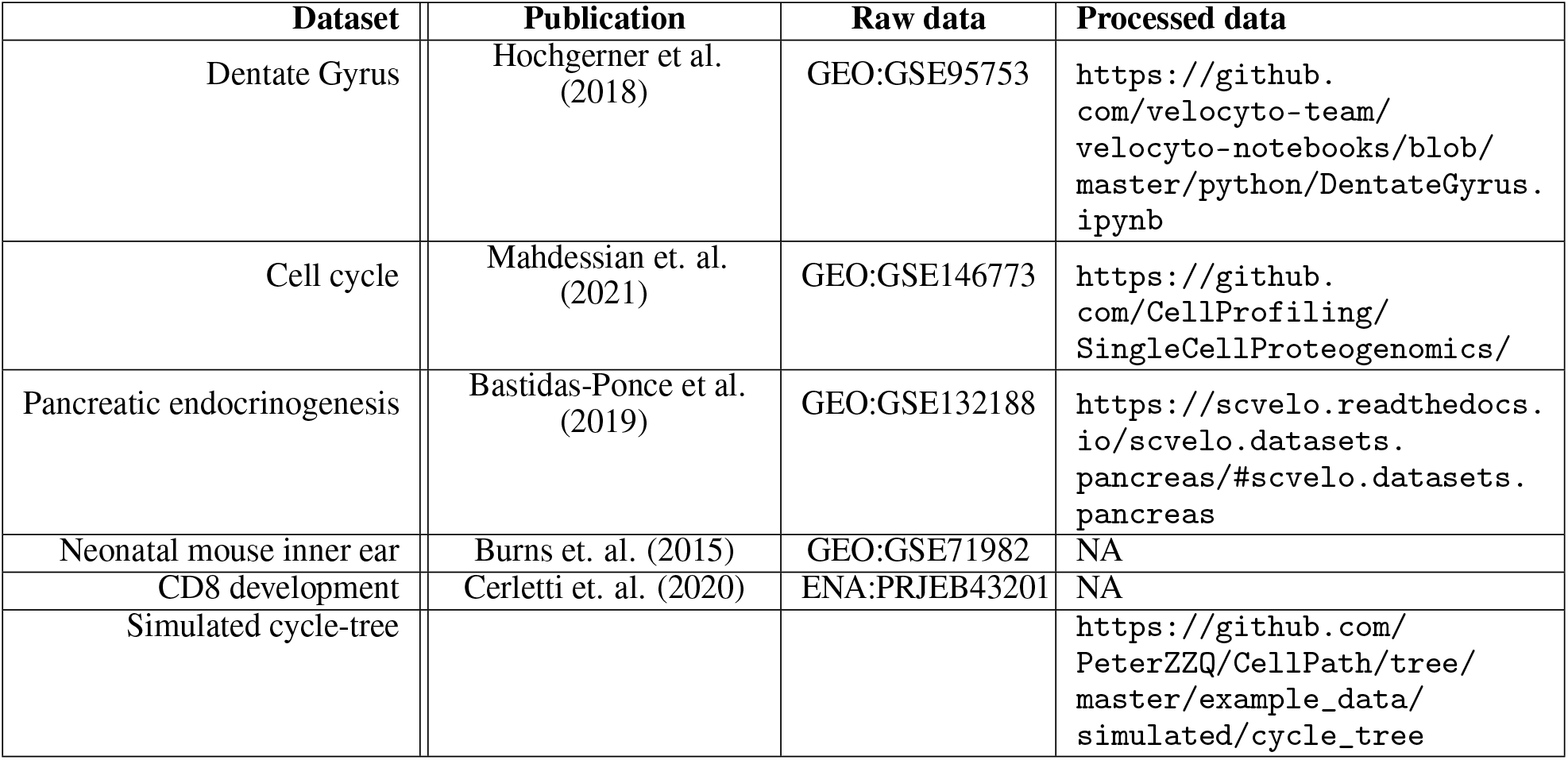
Dataset availability

## Supplementary Tables and Figures

**Table S1:** Features of tools modelling differentiation processes from sc-RNA seq data.

|  | Pseudotime inference | Trajectory coordinates | Branch assignment | Cell fate scoring | Coarse celltype graph | Optional root state input | Optional terminal state input | RNA velocity-based | version |
| --- | --- | --- | --- | --- | --- | --- | --- | --- | --- |
| Cytopath | ✓ | ✓ | ✓ | ✓ | ✓ | ✓ | ✓ | ✓ | v0.1.8.post1 |
| Slingshot | ✓ | ✓ | ✓ |  | ✓ | ✓ | ✓ |  | v2.2.0 |
| Monocle3 | ✓ | ✓ |  |  |  | ✓ |  |  | v1.0.0 |
| VeTra | ✓ |  | ✓ |  |  |  |  | ✓ | 63e638c |
| Cellpath | ✓ | ✓ | ✓ |  |  |  |  | ✓ | v0.2.dev0 |
| Cellrank |  |  |  | ✓ |  |  |  | ✓ | v1.5.1 |
| Scvelo-latent time | ✓ |  |  |  |  | ✓ | ✓ | ✓ | v0.2.4 |
| Directed PAGA |  |  |  |  | ✓ | ✓ | ✓ | ✓ | *scvelo* |

**Figure S1:**
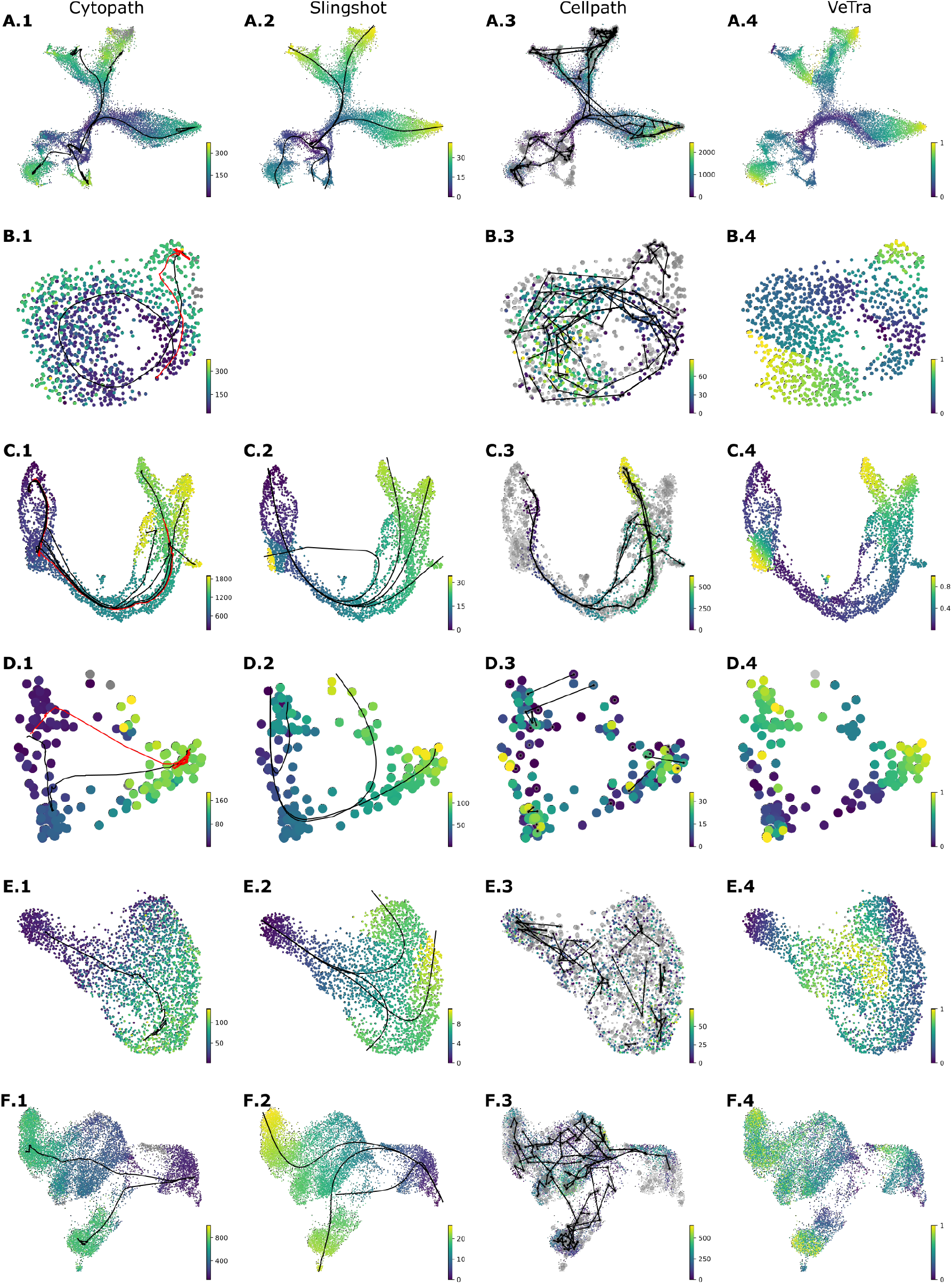
Average pseudotime and trajectories inferred by trajectory inference methods that assign cells to lineages and therefore compute a trajectory specific pseudotime. (A-F) Average pseudotime by Cytopath, Slingshot, Cellpath and VeTra for (A) Dentate Gyrus, (B) cell cycle, (C) pancreatic endocrinogenesis, (D) mouse inner ear (E) neuronal activation and (F) CD8+ T cell exhaustion datasets. Expected behaviour is that all terminal cell states have the highest pseudotime and root states have the lowest pseudotime. Erroneous trajectories initialised at terminal or intermediate states will lead to a patch-like appearance.

**Figure S2:**
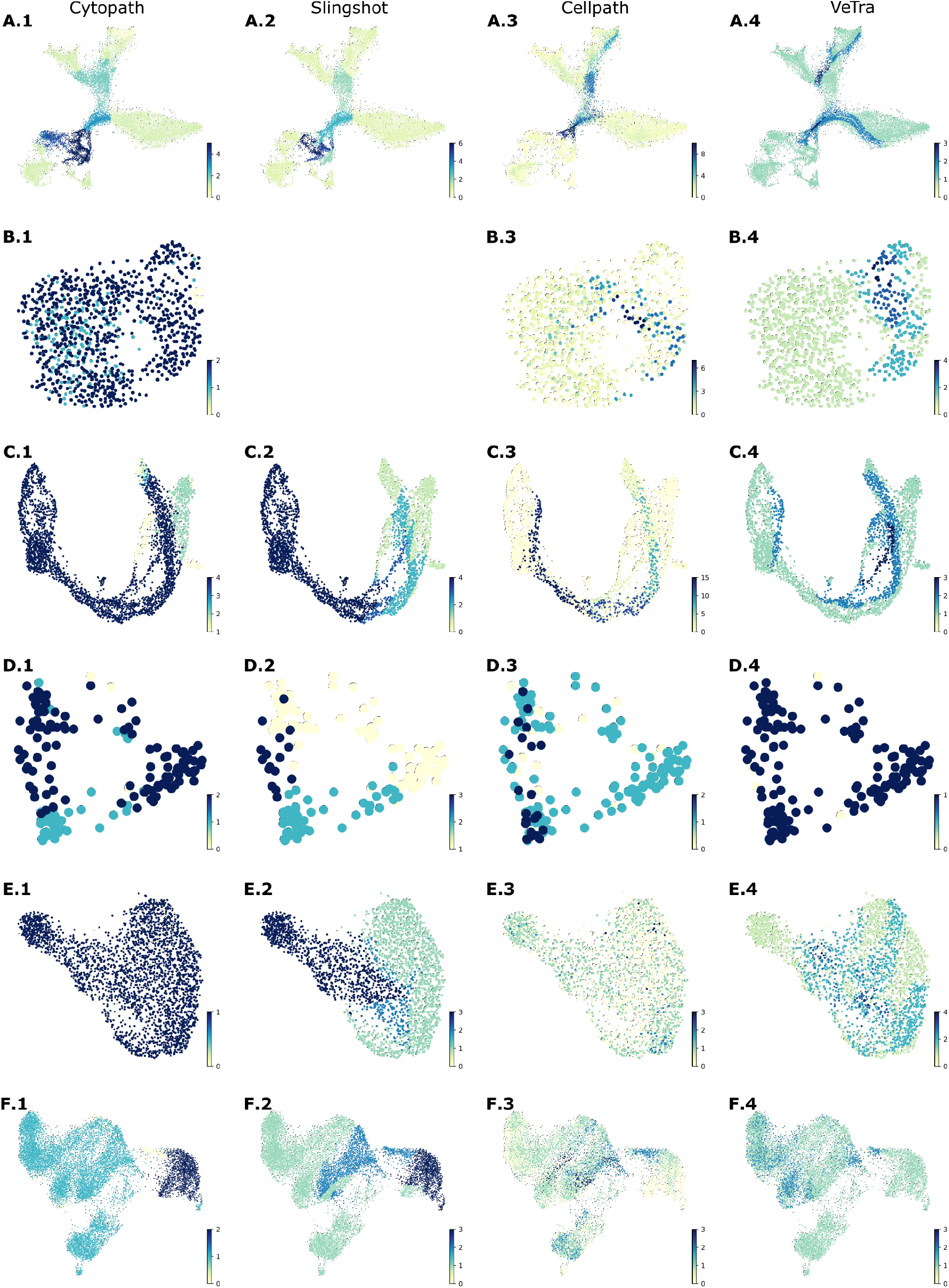
Total number of trajectories to which each cell is assigned by trajectory inference methods. (A-F) Count of trajectories to which each cell is assigned by Cytopath, Slingshot, Cellpath and VeTra for (A) Dentate Gyrus, (B) cell cycle, (C) pancreatic endocrinogenesis, (D) mouse inner ear (E) neuronal activation and (F) CD8+ T cell exhaustion datasets. Expected behaviour is that terminal state cells are assigned to only one trajectory in a multi-terminal state dataset. Cells at the root are typically assigned to all trajectories with each branching event lowering the count. Zero count indicates that the cell was missed by the trajectory inference process entirely.

**Figure S3:**
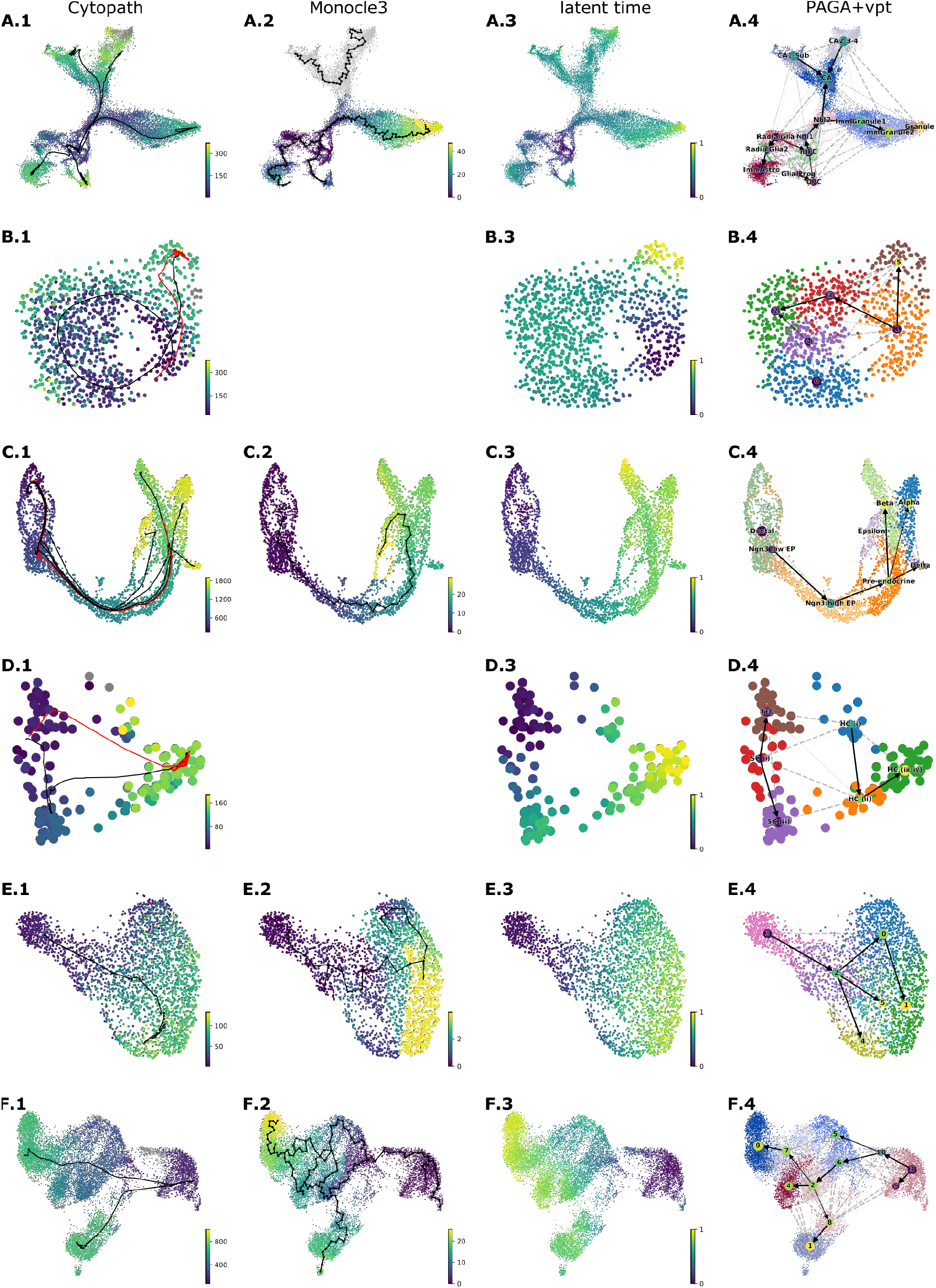
Pseudotime and/or trajectories/graph inferred by methods that compute a global pseudotime per dataset and do not assign cells to lineages. Cytopath pseudotime and trajectories are provided as reference. (A-F) Pseudotime and trajectories/graph by Cytopath, Monocle3, latent time (scvelo) and PAGA with directionality imparted by velocity pseudotime (vpt). (A) Dentate Gyrus, (B) cell cycle, (C) pancreatic endocrinogenesis, (D) mouse inner ear (E) neuronal activation and (F) CD8+ T cell exhaustion datasets.

**Figure S4:**
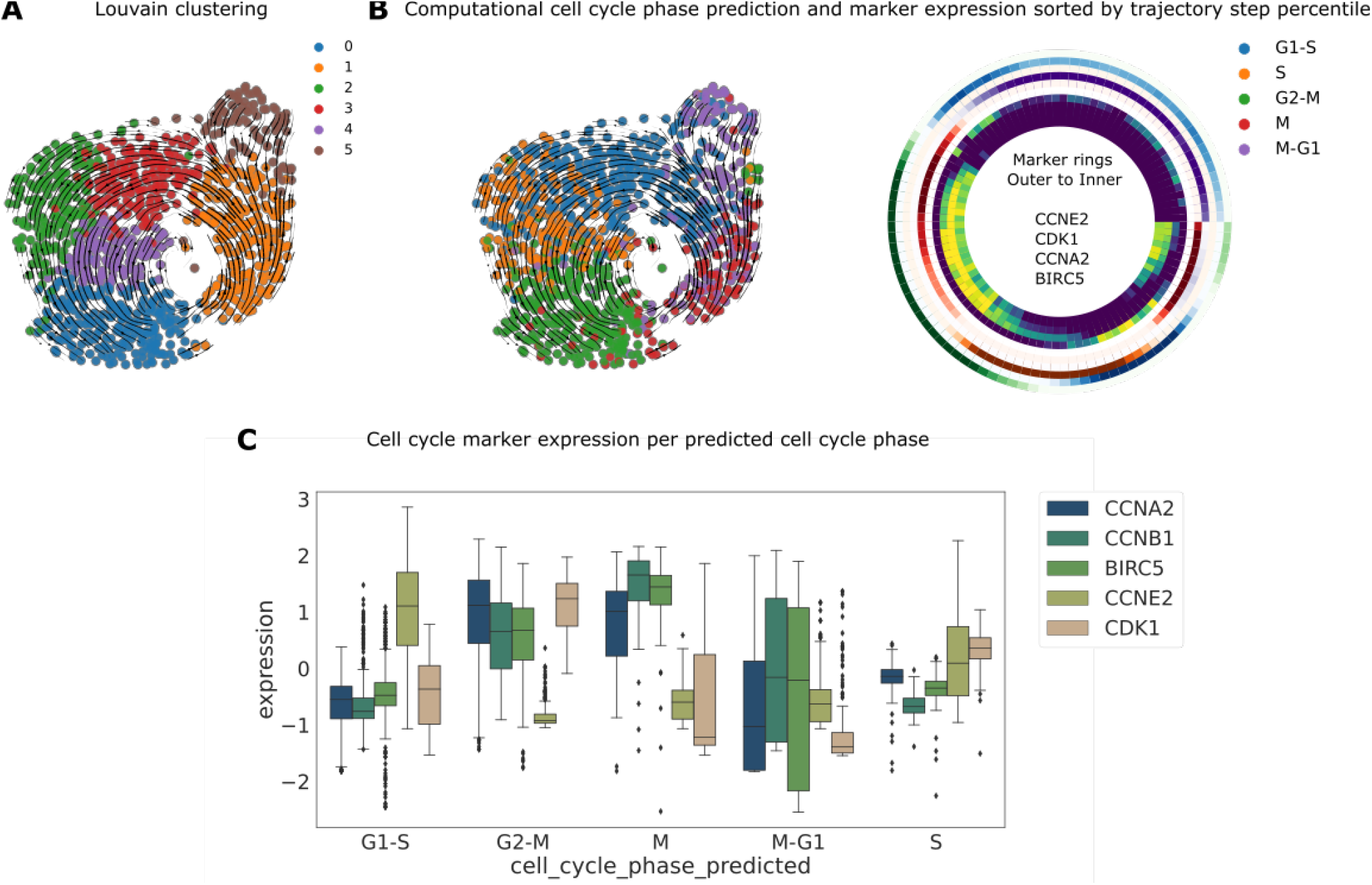
Cell cycle dataset clustering and computational cell cycle phase prediction. (A) Louvain clustering. Cluster 5 corresponds to G1-checkpoint cells. (B) RNA velocity stream plot overlayed on the UMAP projection, annotated with cell cycle phase predicted computationally using expression data. Considering all cell-to-trajectory alignments binned into percentiles, the radial heatmap shows cell cycle phase fraction (outer set of rings) and marker expression (inner set of rings) sorted by trajectory step. The directionality of the radial heatmap is clockwise with the origin at zero degrees. (C) Mean expression of cell cycle markers for each predicted cell cycle phase.

**Figure S5:**
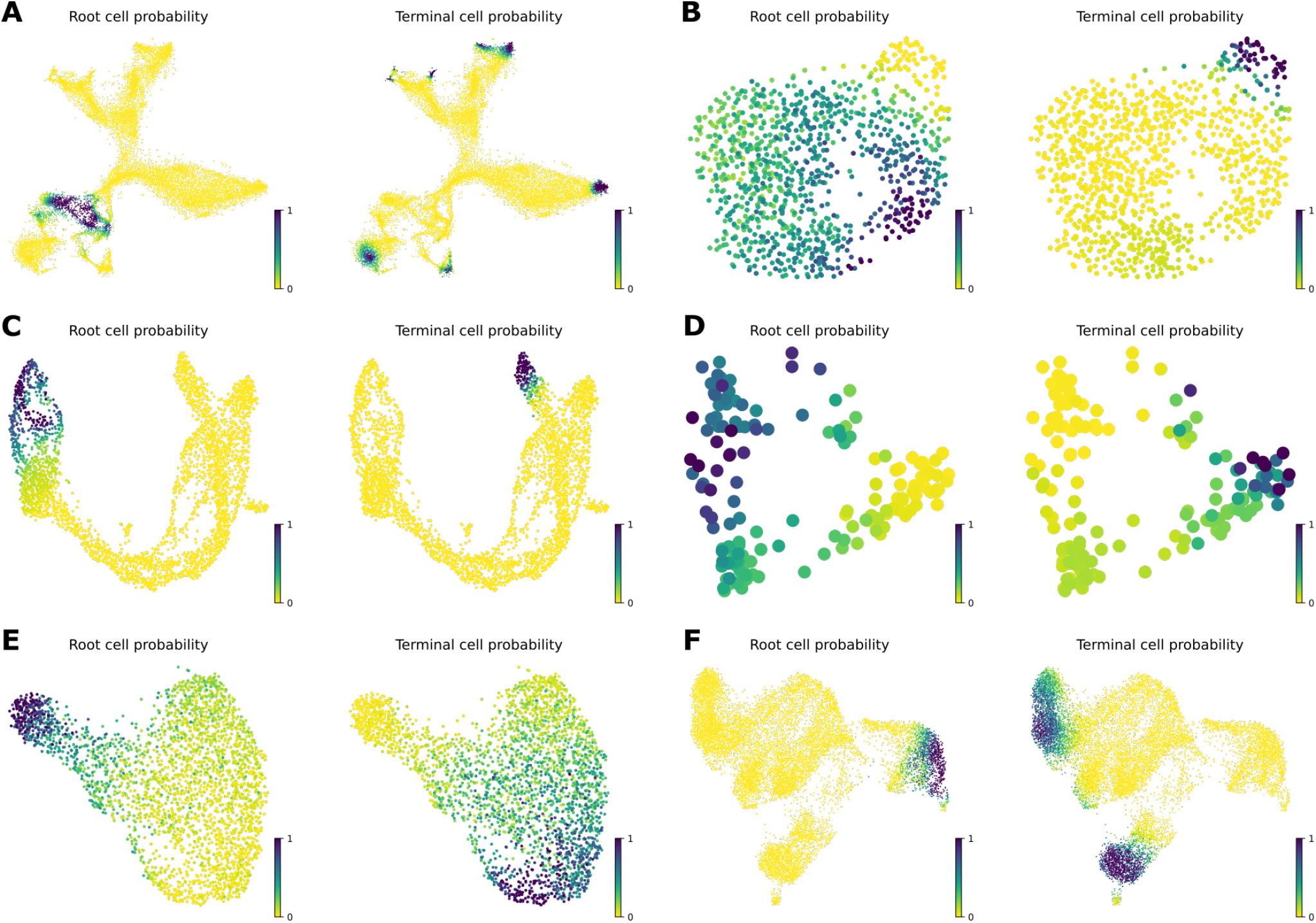
Root and terminal state probability estimation based on RNA velocity. (A-F) Root and terminal state probability of cells belonging to Dentate gyrus (A), cell cycle (B), pancreatic endocrinogenesis (C), neonatal mouse inner ear (D), neuronal activation (E) and CD8 T cell development (F) datasets.

**Figure S6:**
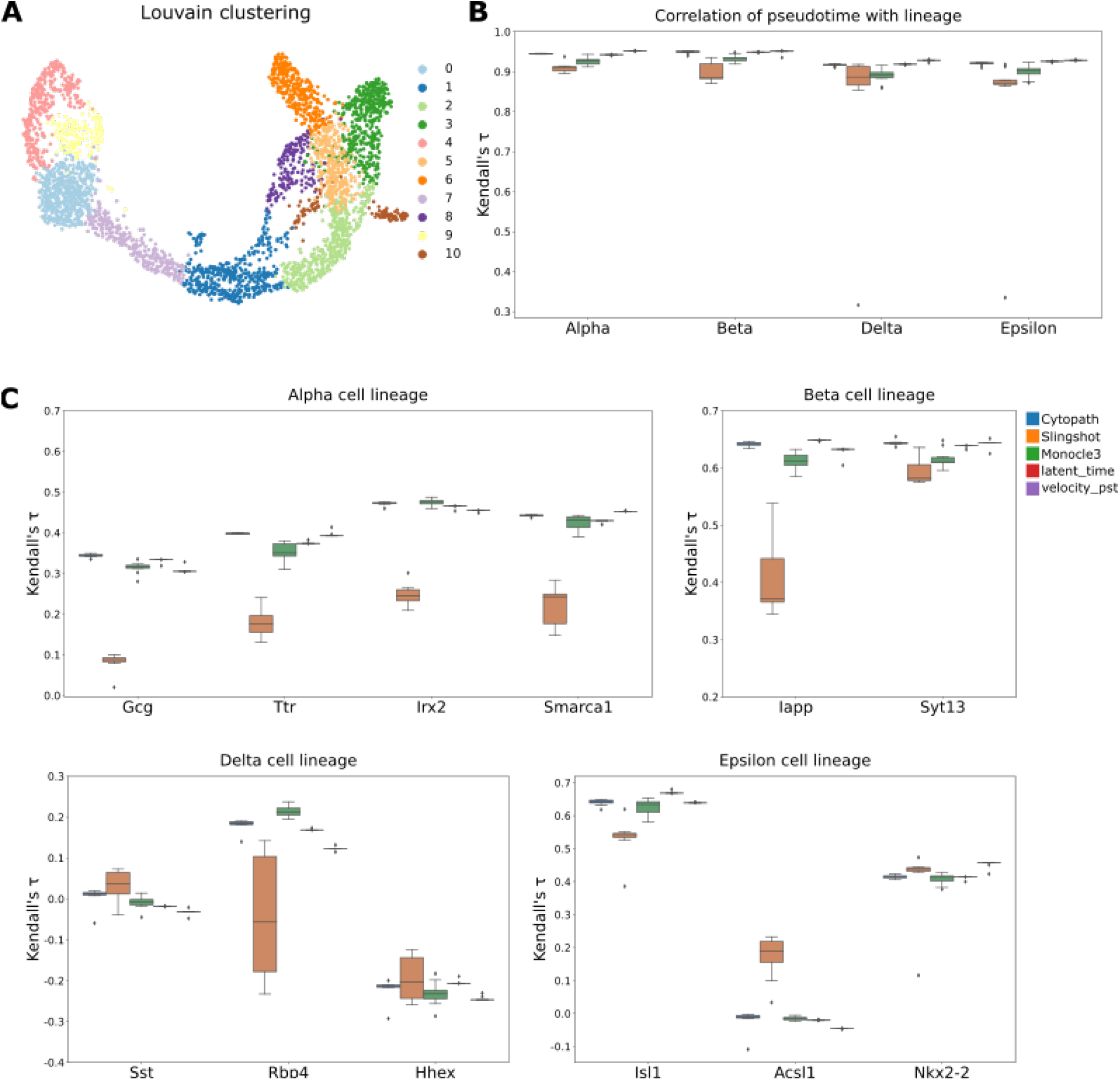
Comparison of methods for the reconstruction of bifurcated differentiation in pancreatic endocrinogenesis. (A) UMAP projection of the pancreatic endocrinogenesis dataset annotated with Louvain clustering. (B) Spearman correlation of pseudotime values assigned by each method to the known ordering of cell types per trajectory. (C) Correlation of pseudotime estimated by each method with markers relevant to the trajectories inferred for each terminal cell type.

**Figure S7:**
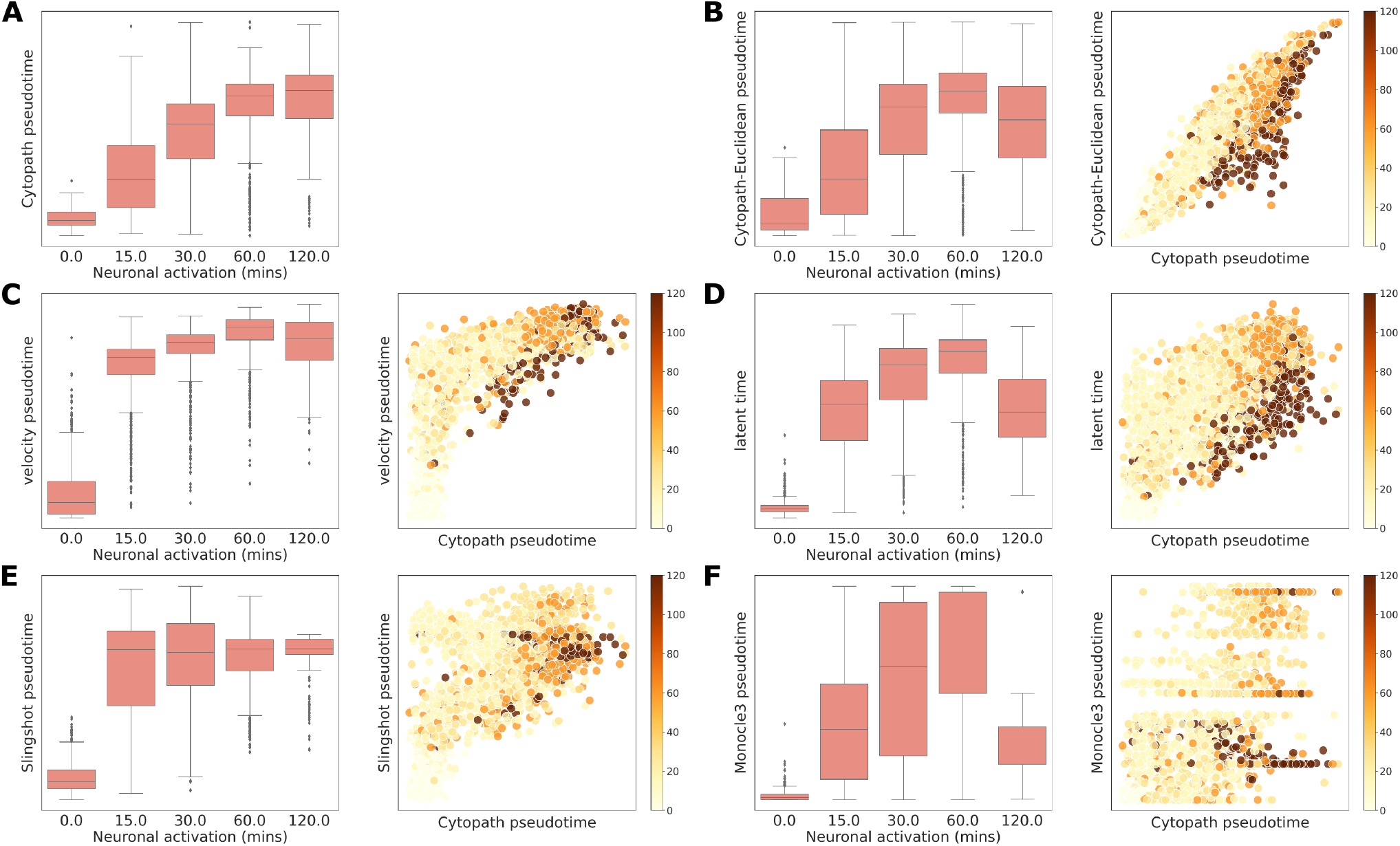
Comparison of pseudotime estimated by each method for the neuronal activation dataset with stimulation time and also with Cytopath pseudotime. (A) Cytopath pseudotime, (B) Cytopath-Euclidean pseudotime (C) velocity pseudotime, (D) latent time, (E) Slingshot pseudotime and (F) Monocle3 pseudotime.

**Figure S8:**
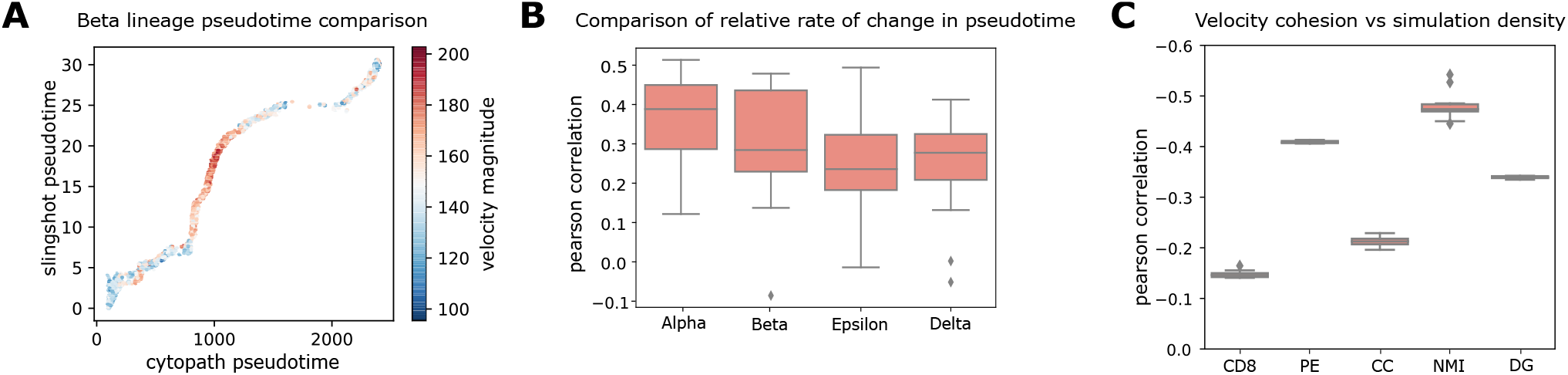
Cytopath pseudotime estimation exhibits dynamic properties of transcriptional time. (A) Slingshot vs Cytopath pseudotime for beta lineage annotated with magnitude of RNA velocity per cell. (B) Relative rate of change of Slingshot pseudotime vs Cytopath pseudotime suggests that Cytopath pseudotime better represents transcriptional dynamics. (C) Correlation of simulation step density for trajectories inferred by Cytopath demonstrate lower density of steps in regions of directed velocity i.e. the trajectories are modelled as transiting faster through regions of consistent directionality and vice-versa.

**Figure S9:**
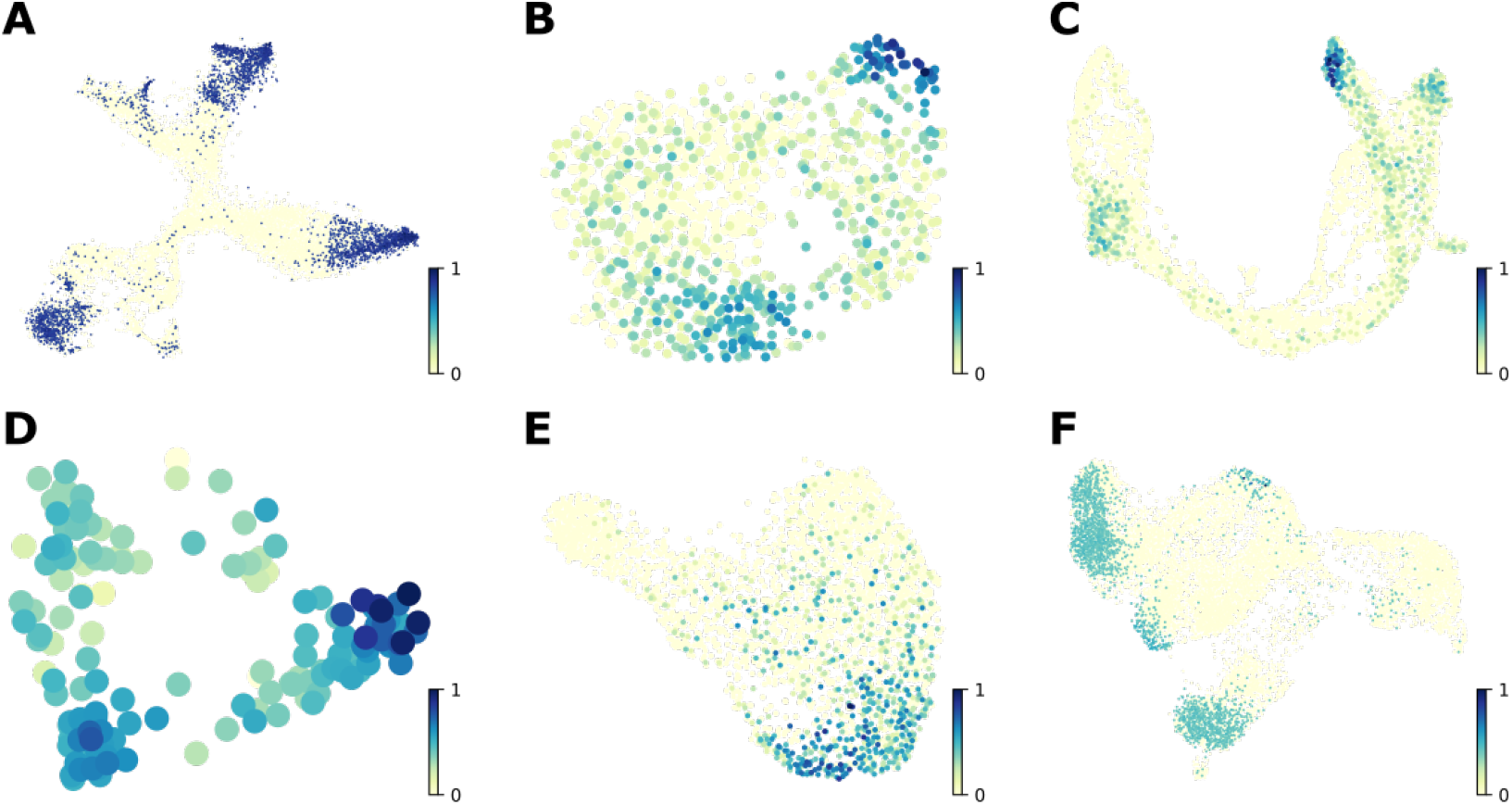
Log of frequency count of undirected simulations (initialised at random cell states) terminating at a cell. Log terminal state frequency per cell scaled to range [0,1] for (A) Dentate gyrus, (B) cell cycle, (C) Pancreatic endocrinogenesis, (D) neontal mouse inner ear, (E) neuornal activation and (F) CD8 T cell development datasets.

**Figure S10:**
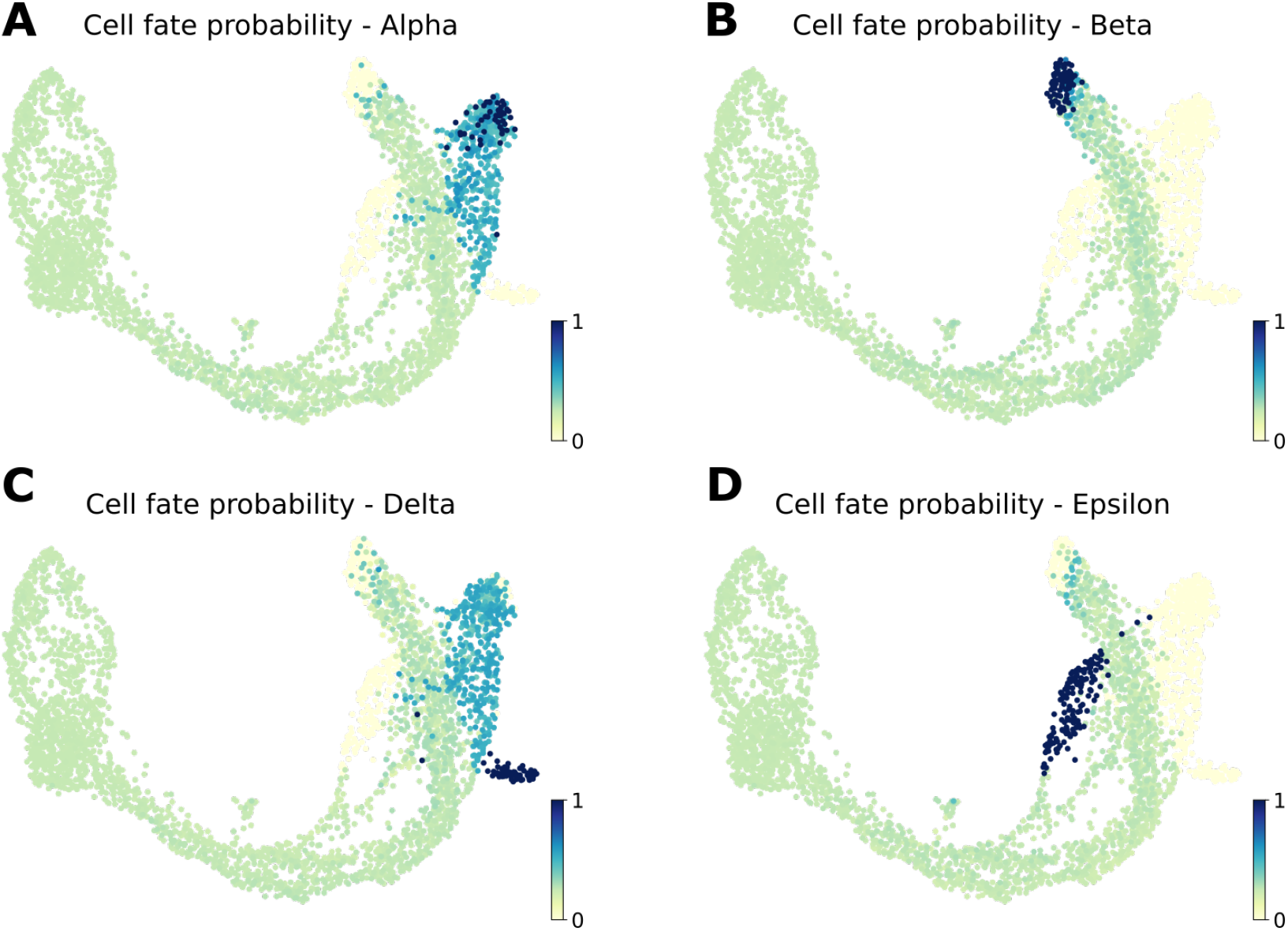
Cytopath estimation of cell fate based on relative alignment of cells. (A-D) Cell fate score per cell of (A) Alpha, (B) Beta, (C) Delta and (D) Epsilon cell type. Cell fate score indicates the likelihood of a lineage representing the differentiation path of the cell.

**Figure S11:**
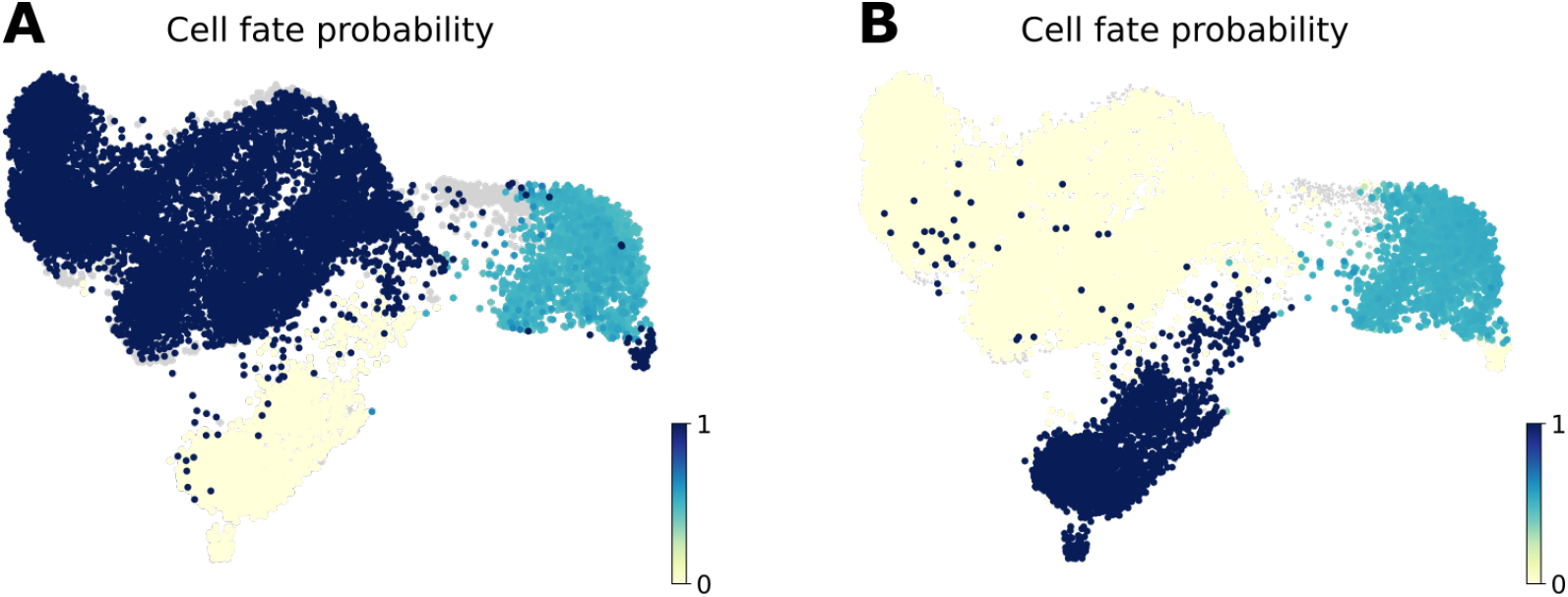
Cytopath estimation of cell fate based on relative alignment of cells. (A-B) Cell fate score per cell of (A) Exhausted CD8+ T cells, (B) Memory-like CD8+ T cells. Cell fate score indicates the likelihood of a lineage representing the differentiation path of the cell.

**Figure S12:**
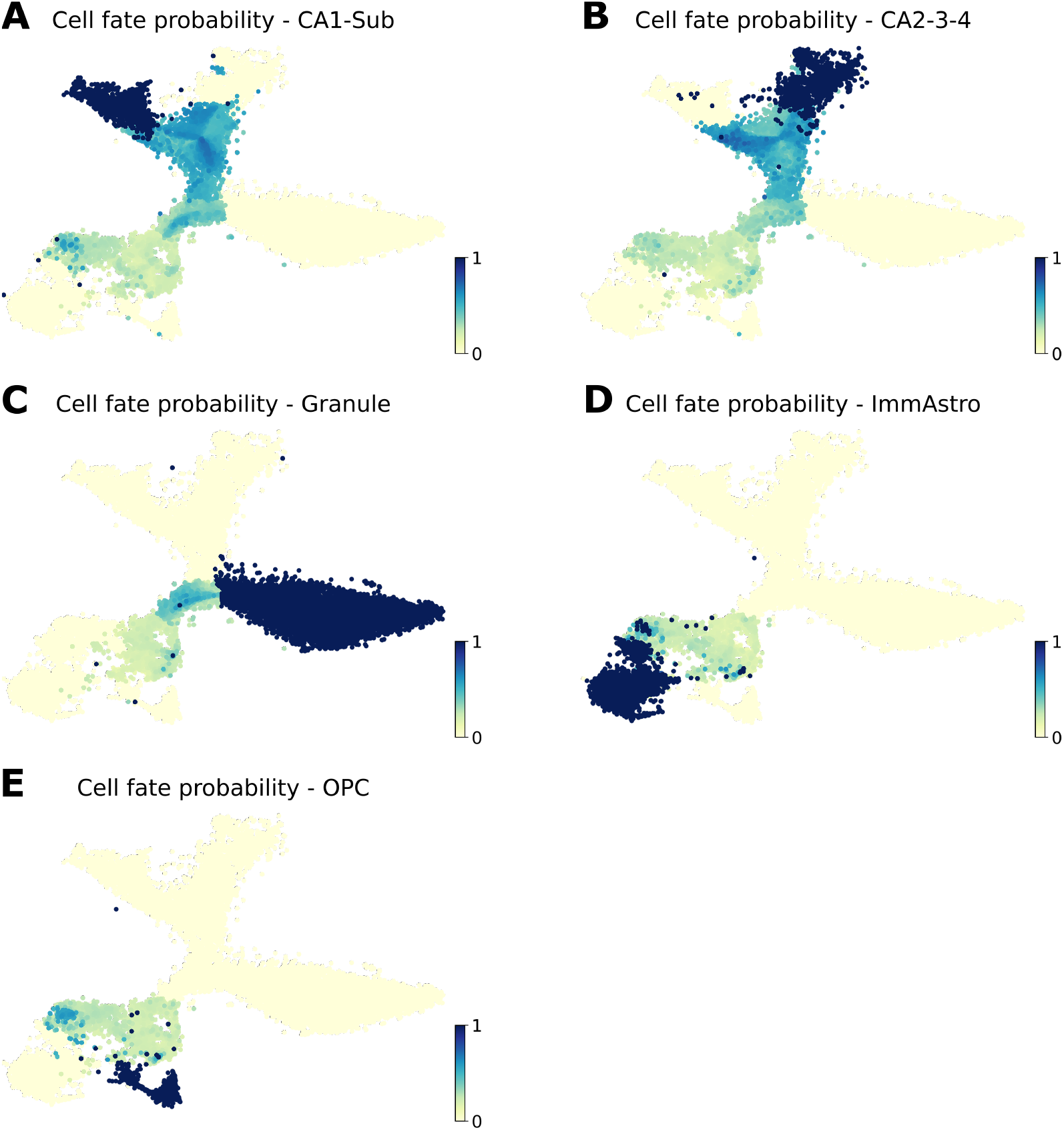
Cytopath estimation of cell fate based on relative alignment of cells. (A-E) Cell fate score per cell of (A) CA1-Subculum, (B) CA2-3-4, (C) Granulocytes, (D) Astrocytes and (E) OPC cell type. Cell fate score indicates the likelihood of a lineage representing the differentiation path of the cell.

**Figure S13:**
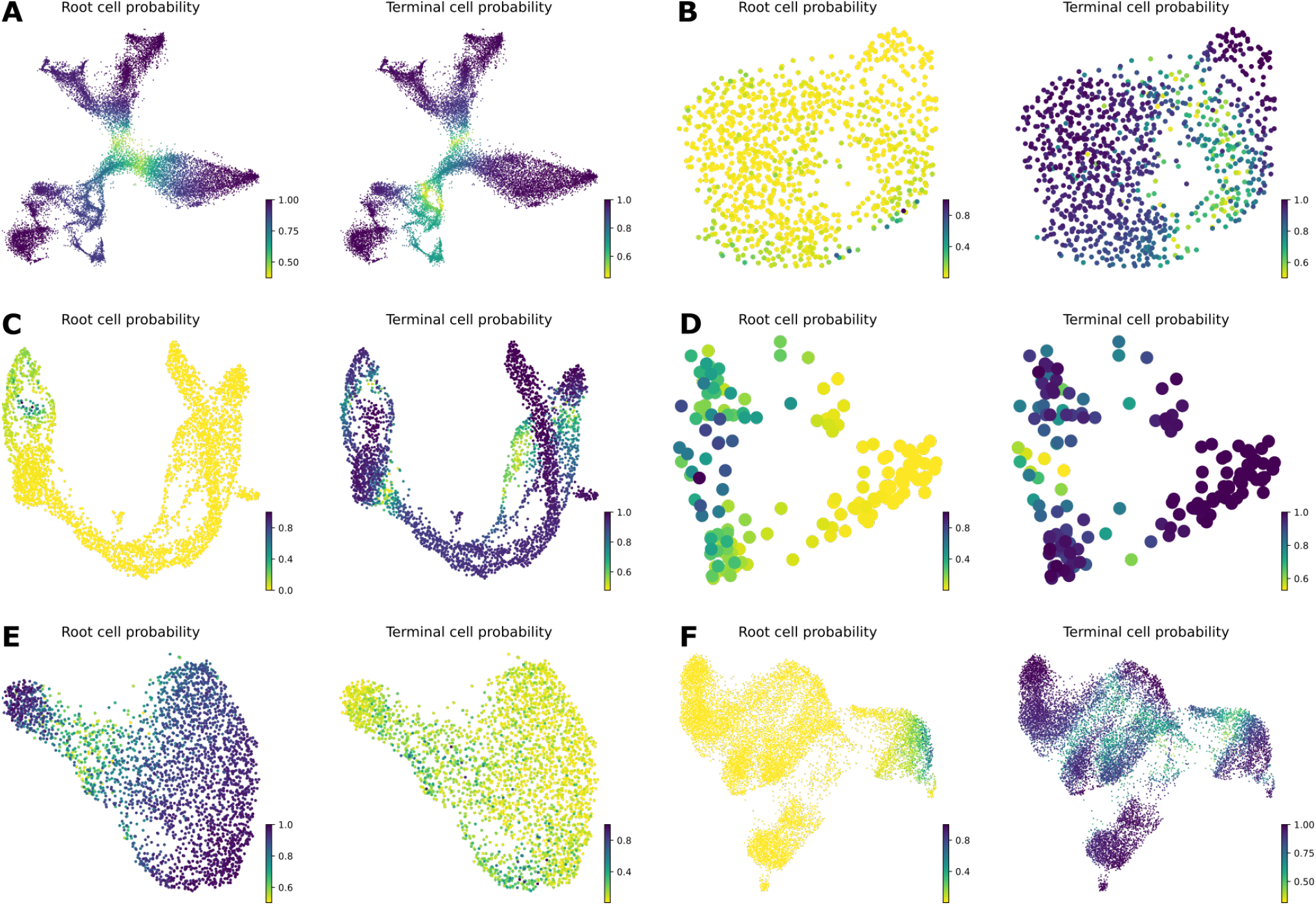
Root and terminal state probability estimation by Cellrank. (A-F) Root and terminal state probability of cells belonging to Dentate gyrus (A), cell cycle (B), pancreatic endocrinogenesis (C), neonatal mouse inner ear (D), neuronal activation (E) and CD8 T cell development (F) datasets.

**Figure S14:**
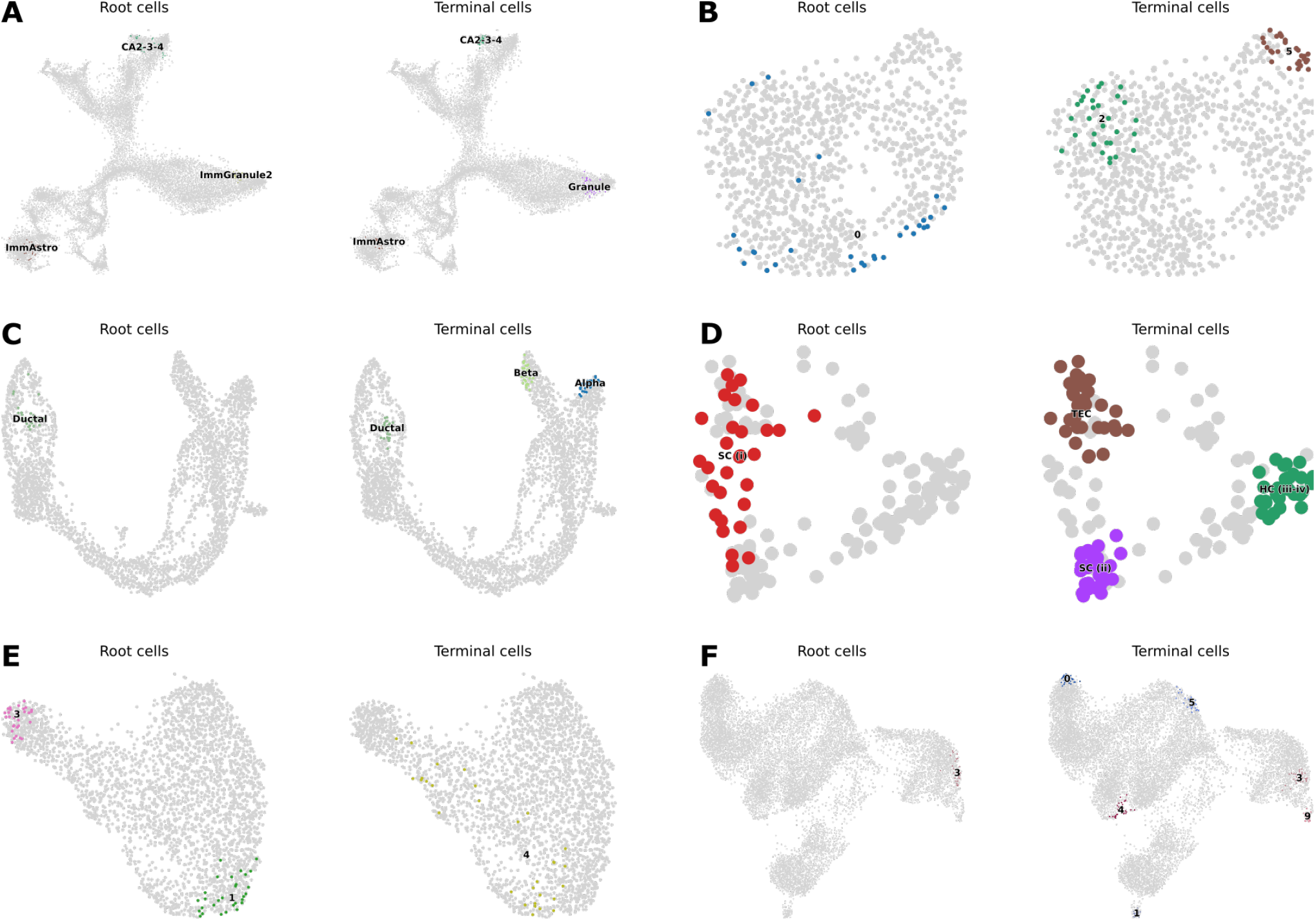
Root and terminal state selection by Cellrank. (A-F) Root and terminal state cells identified by Cellrank in the Dentate gyrus (A), cell cycle (B), pancreatic endocrinogenesis (C), neonatal mouse inner ear (D), neuronal activation (E) and CD8 T cell development (F) datasets.

**Figure S15:**
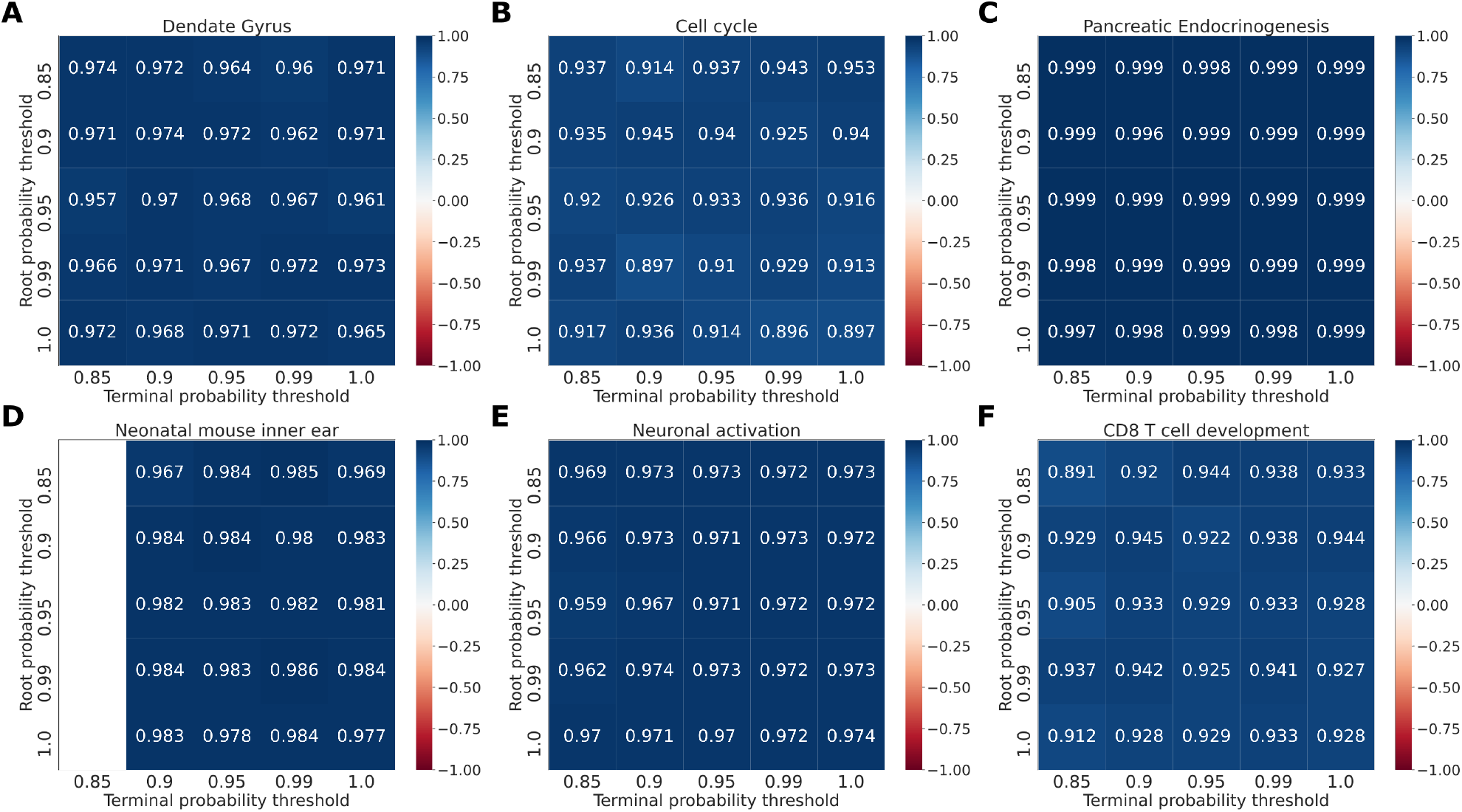
Median Pearson correlation between all pseudotime(s) inferred by Cytopath for the shown thresholds of root and terminal probability while keeping all other parameters at default values. White space indicates that the simulation process failed for that threshold. (A) Dentate Gyrus (B) Cell cycle (C) Pancreatic endocrinogenesis. Note that with default *scvelo* based terimnal state selection only the trajectory to Beta terminal state is inferred. (D) Neonatal mouse inner ear (E) CD8 T cell development.

**Figure S16:**
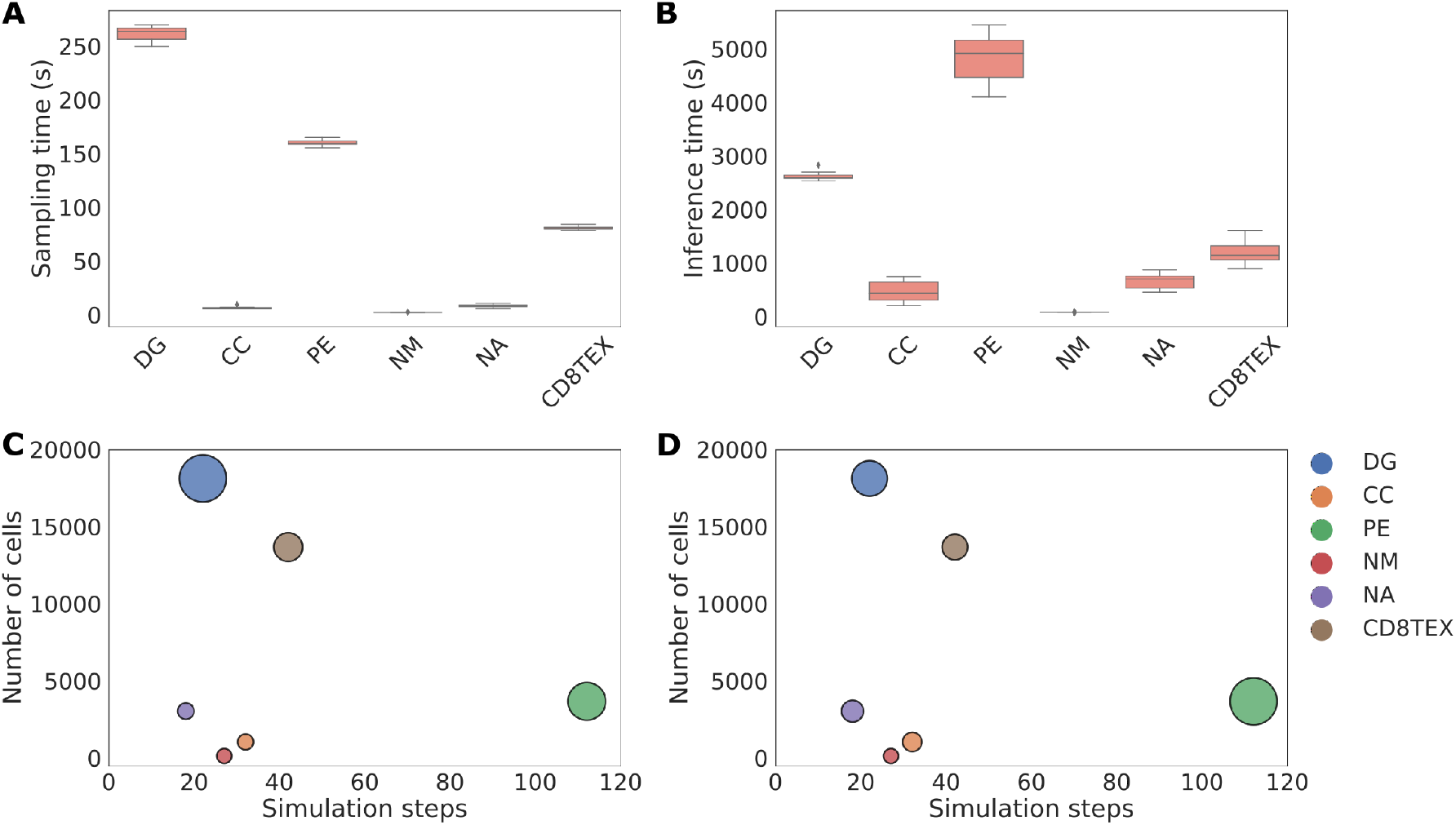
Dependence of Markov sampling and trajectiry inference time on number of simulation steps and size of the dataeset. (A) Time in seconds for Markov sampling per run per dataset, (B) Time in seconds for trajectory and pseudotime inference per run per dataset, (C) Dependence of median Markov sampling time per dataset (relative size of bubble; see values in A) and (D) Dependence of median trajectory and pseudotime inference time per dataset (relative size of bubble; see values in B) for Dentate Gyrus (DG), cell cycle (CC), Pancreatic endocrinogenesis (PE), neonatal mouse inner ear (NM), neuronal activation (NA) and CD8 T cell development (CD8TEX) datasets.

**Figure S17:**
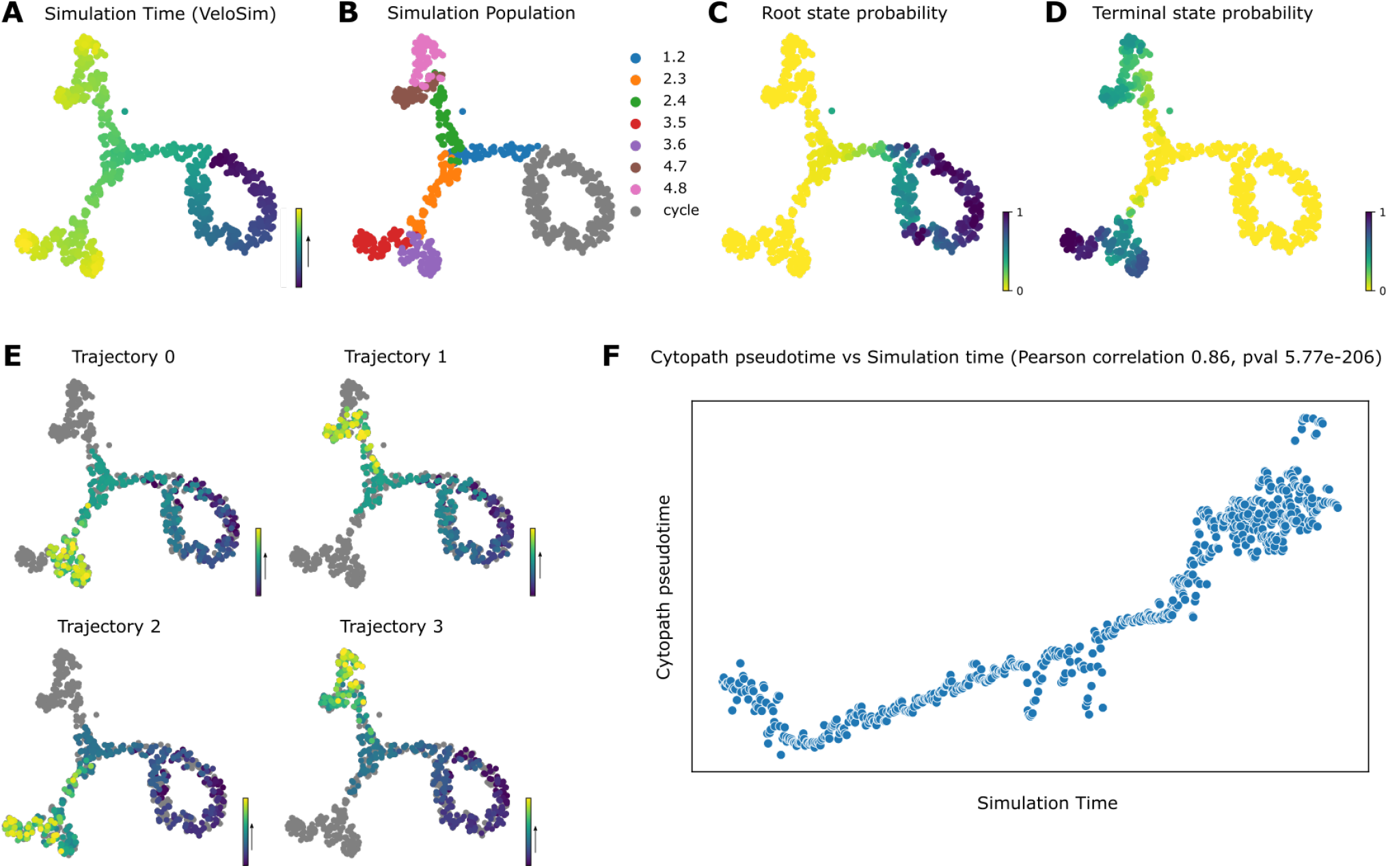
Mean and standard deviation of alignment score per trajectory step. (A) Simulation time from VeloSim. (B) Simulated cell populations from VeloSim. (C-D) Probability estimated using scvelo of cells being (C) root states, (D) terminal states (E) Trajectories inferred using Cytopath overlayed on UMAP annotated with pseudotime. (F) Correlation between Cytopath pseudotime and Simulated time.

## Notes

https://github.com/aron0093/cytopath

### Competing Interest Statement

The authors have declared no competing interest.

### Summary of Updates

Automatic technical parameter selection (Cytopath) has been implemented and results updated to use current package versions. Analysis of pseudotime inferred by Cytopath with respect to internal clock (process time) of cells has been added.

